# STING is a key driver of Japanese encephalitis virus induced inflammatory response

**DOI:** 10.1101/2025.06.11.659027

**Authors:** Simran Chhabra, Dhruvin Patel, Kiran Bala Sharma, Vishal Sah, Krishnan H Harshan, Santosh Chauhan, Manjula Kalia

**Affiliations:** Regional Centre for Biotechnology, NCR Biotech Science Cluster, Faridabad, 121001, India; CSIR-Centre for Cellular and Molecular Biology, Hyderabad, India 500007; Academy for Scientific and Innovative Research (AcSIR), Ghaziabad-201002, India

**Keywords:** cGAS, Flavivirus, Gasdermin D, inflammasome, interferon, MAVS

## Abstract

Interferon (IFN) and inflammation are the key early defence mechanisms that combat pathogen infection. The cytosolic DNA sensor cGAS activates immune signaling via the stimulator of interferon genes (STING) protein. Emerging evidence suggests crosstalk between innate immune DNA and RNA sensing, implicating a role of STING protein in RNA virus infection. This study characterizes STING in the context of Japanese encephalitis virus (JEV), an RNA virus of the *flaviviridae* family. We observe that activation of type I IFN through MAVS is essential for cGAS and STING activation. Knockdown, null mutant and inhibitor studies confirmed that STING restricts JEV replication independently of IFNβ signaling and autophagy. Transcriptomic analysis of STING^gt/gt^ bone-marrow derived macrophages (BMDMs) showed enhanced IFN response, but reduced activation of inflammatory cytokines and chemokines. Phosphorylated STING was recruited on the virus replication complex (RC), marked by the non-structural protein NS1, subsequently triggering the assembly of the NLRP3 inflammasome on the RC. STING proton channel activity was essential for NLRP3 inflammasome activation, IL-1β production, and activation of pyroptotic cell death markers. Sting^gt/gt^ mice, showed higher viremia, earlier disease onset, reduced survival, and decreased brain inflammation. These findings establish STING as a key regulator of JEV-induced inflammation and antiviral defence.

## Introduction

Japanese encephalitis virus (JEV) is an *Orthoflavivirus* transmitted by the bite of an infected *Culex* mosquito. It is the leading global cause of virus induced encephalitis, endemic in east and south-east Asian countries. JEV is neurotropic and the paediatric population is most severely affected with symptoms ranging from mild fever to subsequent severe encephalitis and death [1, 2]. Although vaccines against the virus are available, spread of the disease into previously unaffected geographical areas has been observed. Considering, no effective therapeutics are available, it is crucial to understand the interactions between JEV and host for the development of novel effective antivirals [2].

The host innate immune response is the primary line of defence against virus infection. Its activation depends on the ability of host encoded pathogen recognition receptors (PRRs) to recognize various pathogen associated molecular patterns (PAMPs) or host derived damage-associated molecular patterns (DAMPs) [3]. The host genome encodes for different classes of PRRs including cell surface and endosomal toll-like receptors (TLRs), nucleotide oligomerization domain (NOD)-like receptors (NLRs also called NACHT, LRR and PYD domain proteins), cytosolic nucleic acid sensors like the retinoic acid-inducible gene I (RIG-I)-like receptors (RLRs), cyclic guanosine monophosphate-adenosine monophosphate (cGAMP) synthase (cGAS) and C-type lectin receptors (CLRs) [4]. Upon recognition of PAMPs, these PRRs induce a cascade of signaling events resulting in the production of type I interferon (IFN) thereby generating an antiviral state inside the cell [5]. Studies have shown that RIG-I, TLR2, TLR3, TLR7, and TLR8 play key roles in modulating JEV-induced immune and inflammatory responses in microglia, astrocytes as well as neuronal cells [6–8]. Our earlier study has reported the activation of cyclic guanosine monophosphate-adenosine monophosphate (cGAMP) synthase (cGAS)-stimulator of interferon genes (STING) pathway during JEV infection and a crucial antiviral role of cGAS protein in JEV-infected fibroblasts [8].

The adaptor protein STING is an endoplasmic reticulum (ER)-associated membrane protein with diverse functions. It is best known for its role in sensing intracellular DNA through cGAS-STING pathway, leading to the activation of IFN and inflammatory responses [9, 10]. Additionally, non-canonical functions of STING have recently gained considerable attention. STING can modulate autophagy [11], senescence [12], inflammasome [13–16], protein translation [17], cellular condensation [18], metabolic pathways [19], lysosome recovery and biogenesis [20, 21] as well as cell death [22, 23]. In context of virus infection, type I IFN and inflammatory cytokines are generally considered the principal downstream effectors of STING-mediated antiviral responses [24–26]. However, STING also restricts virus replication through IFN independent pathways including autophagy induction [27, 28], translation inhibition [29], cell death [30, 31] and inflammasome activation [13].

While, the protective role of STING in response to DNA virus infection is well described, the mechanisms underlying its involvement in RNA virus infection remain to be elucidated. Although RNA and DNA sensing pathways are traditionally distinguished based on their roles in IFN expression, crosstalk between STING and RNA-sensing RIG-I-Mitochondrial anti-viral signaling protein (MAVS) pathway has also been suggested in studies involving HIV-1 [32], and dengue virus (DENV) [33]. The critical role of STING in innate immune defense against RNA viruses is further underscored by the identification of RNA viruses that actively antagonize STING-dependent antiviral response [34, 35]. Flaviviruses, particularly have developed unique strategies to modulate STING signaling [36, 37].

In this study, we have characterized the role of STING during JEV infection *in vitro* and in mouse model of disease. We observe that the expression of cGAS, and STING is regulated by MAVS-IFN signaling pathway. STING restricted virus replication in diverse cell types: fibroblasts, neuronal cells and bone-marrow derived macrophages (BMDMs). The antiviral role of STING was found to be independent of interferon response and canonical autophagy machinery. Transcriptomic analysis of JEV infected STING^gt/gt^ BMDMs showed significant inhibition of non-canonical inflammasome activation resulting in reduced secretion of pro-inflammatory cytokines and chemokines. We also found that proton channel activity of STING was critical for activation of NLRP3 inflammasome as well as cytokine release. Interestingly, pSTING also localized to the viral replication complex (RC), marked by NS1, facilitating NLRP3 inflammasome assembly. Finally, in STING^gt/gt^ mice, we observed higher viremia, earlier disease onset, reduced survival, and decreased brain inflammation. Our study establishes STING as a key regulator of JEV-induced inflammation.

## Results

### JEV infection activates STING which localises to virus replication membranes

Our previous study demonstrated that JEV infection results in activation of cGAS and STING [8]. Here, we first investigated the expression pattern of innate immune effector proteins and observed enhanced phosphorylation of STING, TBK1 and IRF3, and increased levels of cGAS in JEV infected mouse embryonic fibroblasts (MEFs) (Fig. 1a). We further confirmed this observation by assessing the phosphorylation status of STING and TBK1 over a JEV-infection time-course (Fig. S1a-c). STING is primarily localized in the ER, where it plays a central role in regulating innate immune signaling [10]. We also evaluated STING and pSTING localization through immunofluorescence, and observed punctate and scattered distribution in mock-infected cells, which changed to well-defined clusters in virus-infected cells (Fig 1b, S1d). Interestingly, both STING and pSTING showed remarkable colocalization with the JEV-NS1 protein that marks the virus replication complex [38–40]. JEV replication membranes are derived from the ER, and are enriched in several ER-associated degradation (ERAD) effector proteins [39] and non-lipidated LC3 (LC3-I) [41]. These data suggest that JEV infection activates STING which is recruited to the sites of virus replication.

**Figure 1:**
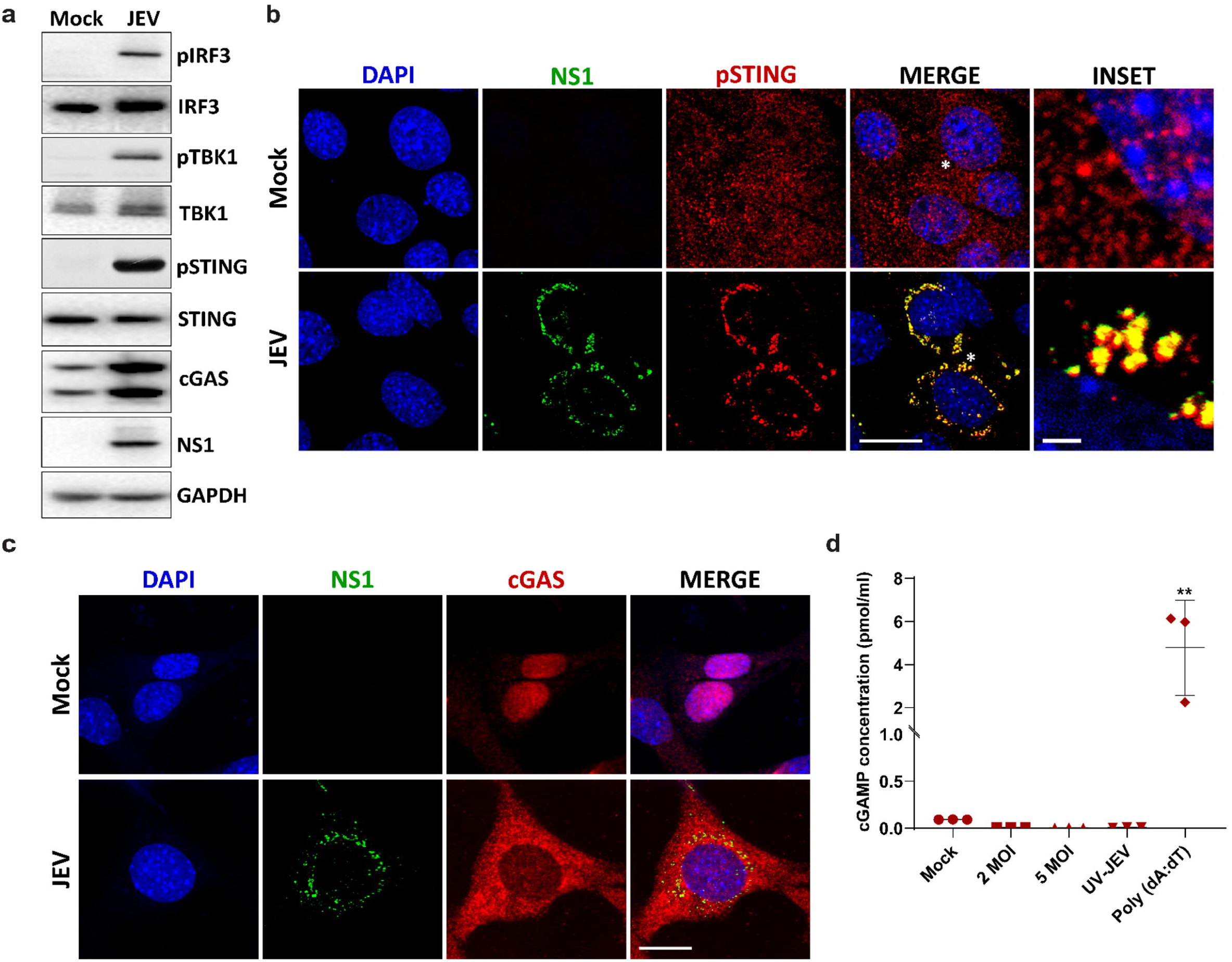
JEV infection activates STING which localises to virus replication membranes. MEFs were mock/JEV infected at 2 MOI for 24 h. (a) Protein lysates were immunoblotted for pIRF3, IRF3, pTBK1, TBK1, cGAS, pSTING, STING, JEV-NS1 (infection control) and GAPDH (loading control). MEFs were mock/JEV infected with JEV (5 MOI, 24h). Cells were stained with (b) JEV-NS1 (green) and pSTING (red) antibodies or and (c) with JEV-NS1 (green) and cGAS (red) antibodies. Nucleus was stained with DAPI. Images were acquired on a confocal microscope using a 63x objective. Scale bar 10µm. Images are representative of two independent experiments. (d) MEFs were either mock/JEV/ UV-JEV infected at 2 & 5 MOI for 24 h or transfected with Poly (dA:dT) at 1 μg/ml and harvested at 12 h. Cell lysates were prepared and cGAMP production was assessed using DetectX® 2ʹ,3ʹ-Cyclic GAMP ELISA Kit.

Since cGAS upregulation was also observed in JEV-infected cells, we checked its subcellular localization. The DNA-sensing function and downstream activation of innate immune response via cGAS depends on its localization in cytosol [42]. Immunofluorescence and subcellular fractionation assays of virus-infected cells showed both upregulation and enrichment of cGAS in the cytosol, and a concomitant reduction in its nuclear levels (Fig 1c, S1e). Cytosolic cGAS was diffuse in JEV infected cells and did not show any localization to the virus replication membranes. Further, no cGAMP production was detected upon JEV infection ranging from low to high multiplicity of infections (MOIs) (Fig. 1d), suggesting that activation of cGAS occurs independently of cGAMP production.

### cGAS and STING activation during JEV infection is regulated by MAVS-IFN signaling

We next investigated if there is any crosstalk between the RNA-sensing RIG-I-MAVS signaling and cGAS-STING during JEV infection. For this, we used *Mavs^-/-^* MEFs (Fig. 2a) and observed that these cells were unable to activate type-I IFN during JEV infection (Fig. 2b), resulting in very high virus replication (Fig. 2c, d). We also observed significantly reduced transcript levels of both *cgas* and *Sting* which did not increase upon virus infection (Fig 2e,f). Subsequently, the protein levels of cGAS and STING also remained low and unchanged in virus-infected MAVS deficient cells (Fig 2 g-i). These data suggest that MAVS is crucial for activation of cGAS and STING. To explore the role of IFN signaling, we also tested BMDMs isolated from WT and AG129 mice, a double-knockout mouse model lacking IFN alpha/beta and gamma receptors (IFNαβγR-/-) [43]. Due to lack of IFN signaling (Fig. 2j), virus replication in these cells was very high as compared to WT BMDMs (Fig 2k-l). Consistent with our results in MAVS deficient cells, we found that the mRNA (Fig. 2m,n) and protein levels of cGAS and STING (Fig. 2o-q) were significantly low in JEV infected AG129 BMDMs. These results clearly highlight that activation of MAVS and further IFN signaling is responsible for cGAS and STING enhancement, suggesting that these molecules are likely to function as interferon stimulated genes (ISGs) during JEV infection.

**Figure 2:**
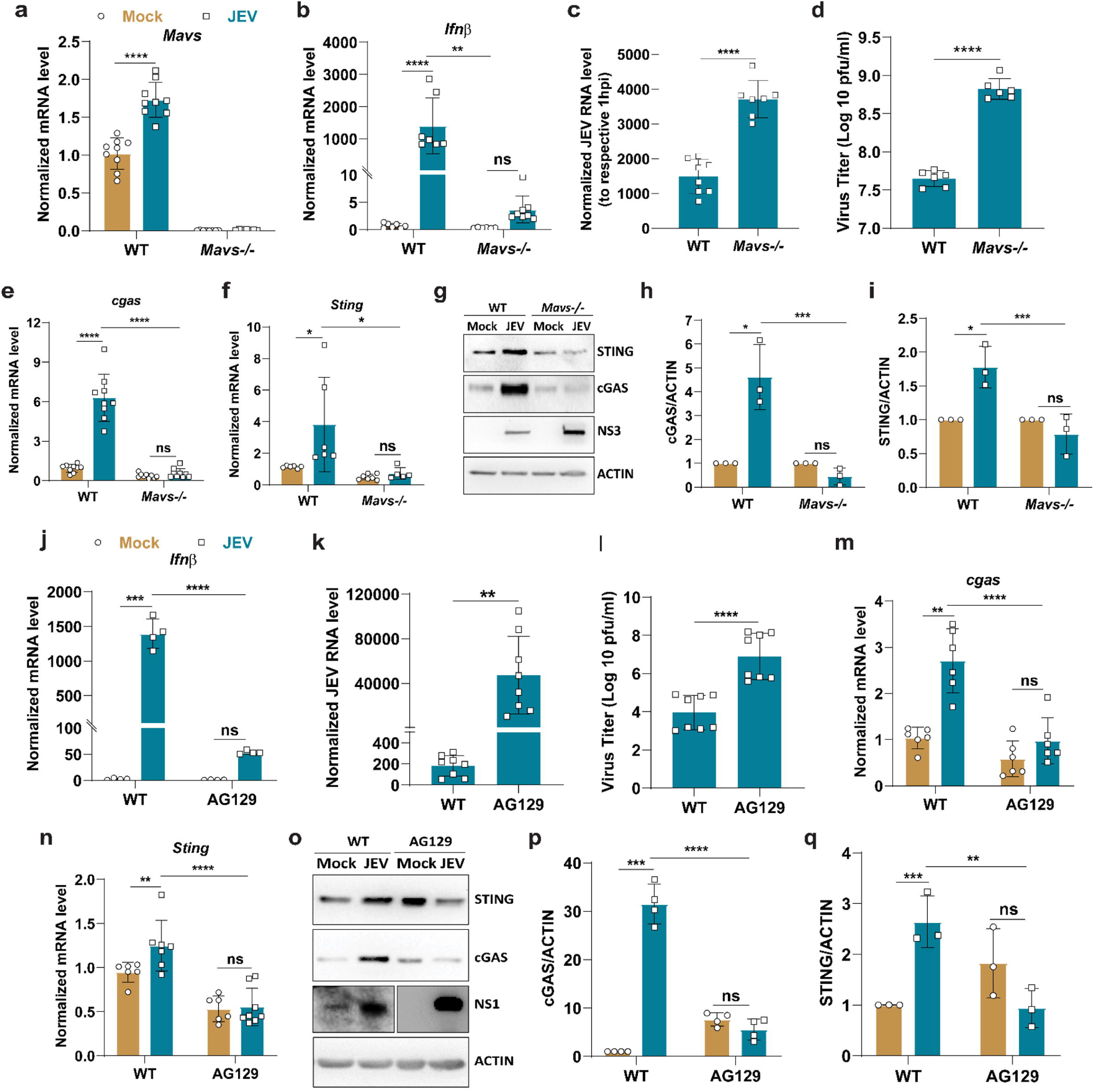
cGAS and STING activation during JEV infection is regulated by MAVS-IFN signaling. MEFs isolated from WT and Mavs^-/-^ C57BL/6 embryo were mock/JEV infected at 2 MOI for 1 h and 24 h. Total RNA was isolated to quantitate mRNA expression of *Mavs, Ifnβ, cgas, Sting* and JEV RNA levels (a,b,c,e,f). (d) Culture supernatant collected was used to determine extracellular virus titers using plaque assays. (g) Protein lysates were immunoblotted with STING, cGAS, JEV-NS3 and ACTIN (loading control) antibodies. (h,i) BAR graph shows the normalized cGAS and STING protein expression (JEV/mock) for three independent experiments. BMDMs from WT and AG-129 C57BL/6 mice were mock/JEV infected at 5 MOI for 24 h. Total RNA was isolated to quantitate JEV-RNA levels (k) and *Ifnβ, cgas and Sting* transcript levels (j,m,n) by qRT-PCR. (l) Culture supernatant collected was used to determine extracellular virus titers using plaque assays. (o) Protein lysates were prepared from mock/JEV infected BMDMs and western blot analysis was done for STING, cGAS, JEV-NS1 and ACTIN (loading control). (p,q) BAR graph shows the normalized cGAS and STING protein expression (JEV/mock). Data presented is mean ± SD of values obtained from 3 independent experiments. Student t-test was used to calculate P values. *P < 0.05, **P < 0.01, ****P < 0.0001. ns, non-significant.

### STING restricts virus replication independent of IFNβ response

To evaluate the role of STING in JEV replication, we utilized STING^gt/gt^ primary MEFs [44]. The Goldenticket (Gt) mutant mouse model harbors a point mutation (T596A) in Sting, causing an isoleucine-to-asparagine substitution (I199N) in the STING protein, which results in the loss of its function and impaired type I IFN signaling. A time-course analysis of virus infection in WT and STING^gt/gt^ JEV infected MEFs showed similar RNA levels till 6 hpi (Fig. 3a). This time-point corresponds to early events of infection leading up to formation of the virus replication factories and generation of double-strand (ds) RNA replicative intermediate [45]. Subsequently significantly enhanced viral RNA levels (Fig. 3a), and titres (Fig. 3b) were observed at 12 and 24 hpi in the STING^gt/gt^ MEFs. Significantly higher levels of viral protein (Fig.3c-d) were also indicative of enhanced virus replication under conditions of STING deficiency. Activation of innate immune signaling was also observed, as seen by increased phosphorylation of TANK-binding kinase 1 (TBK1) and Interferon Regulatory Factor 3 (IRF3), and enhanced transcript and secretory levels of IFNβ in JEV infected STING ^gt/gt^ MEFs compared to WT cells (Fig. 3e-h). However, with pI:C treatment reduced TBK1 and IRF3 phosphorylation, and reduced IFNβ release was observed in STING^gt/gt^ MEFs (Fig. 3e, f, i). These results indicate that while STING^gt/gt^ MEFs are intrinsically defective in producing a type-I IFN response with pI:C treatment, they have no impact on innate immune activation and IFN production in JEV infection. It is likely that higher viremia in STING^gt/gt^ MEFs resulted in significantly higher innate immune activation and IFNβ production.

**Figure 3:**
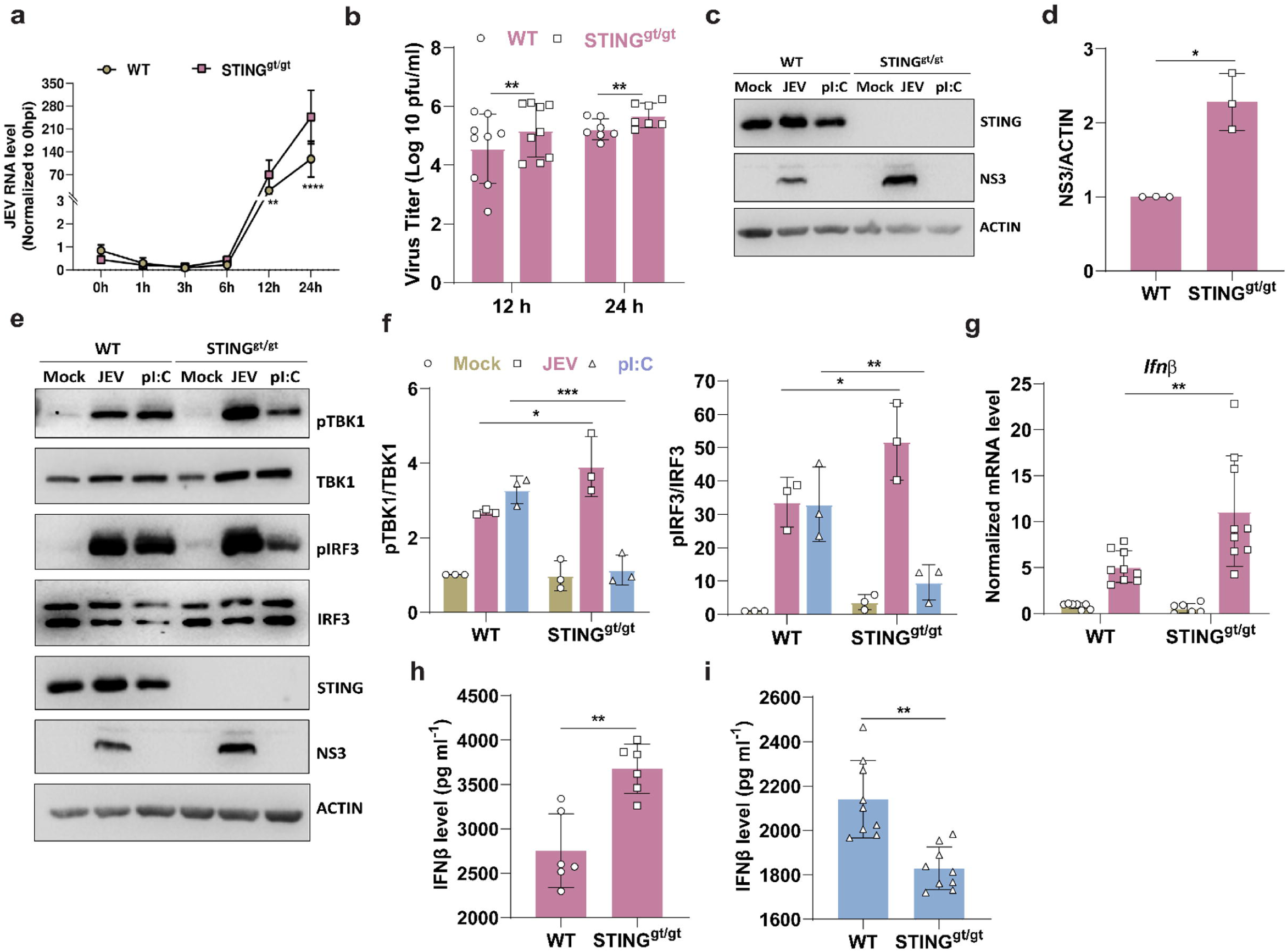
STING restricts virus replication independent of IFNβ response. MEFs isolated from WT and STING^gt/gt^ C57BL/6 13 days embryo were mock/JEV infected at 2 MOI for 0, 1, 3, 6, 12 and 24 h. Total RNA was isolated to quantitate JEV-RNA levels (a). (b) Culture supernatant was collected from 12 and 24 h timepoint to determine extracellular virus titer by plaque assay. (c) Protein lysates from 24 h timepoint were immunoblotted with STING, JEV-NS3 and ACTIN (loading control) antibodies. (d) Bar graph shows the normalized JEV-NS3 protein expression (JEV/mock). MEFs were either mock/JEV infected at 2 MOI for 24 h or transfected with pI:C (1 μg/ml) for 6 h. (e) Protein lysates were prepared and immunoblotted with pTBK1, TBK1, pIRF3, IRF3, STING, JEV-NS3 and ACTIN (loading control) antibodies. (f) Bar graph shows the normalized pTBK1/TBK1 and pIRF3/IRF3 protein expression (JEV/mock) for three independent experiments. Total RNA was isolated from mock/JEV infected MEFs and transcript levels of *Ifnβ* (g) were determined by qRT-PCR. (h) Culture supernatant was collected to determine the extracellular release of IFNβ from mock/JEV infected MEFs. (i) WT and STING^gt/gt^ MEFs were transfected with 1 μg/ml Poly(dA:dT) for 6 h and culture supernatant was collected to determine the extracellular release of IFNβ. Data presented is mean ± SD of values obtained from 3 independent experiments. Student t-test was used to calculate P values. *P < 0.05, **P < 0.01, ****P < 0.0001. ns, non-significant.

We strengthened these observations through siRNA mediated knockdown of STING in MEFs. Depletion of STING (Fig. S2a) resulted in marked enhancement of JEV replication, as seen by significantly increased viral RNA (Fig. S2b), titers (Fig. S2c), and protein expression (Fig. S2d-e). We also observed enhanced phosphorylation of TBK1 and IRF3 (Fig. S2d, f-g), alongside increased IFNβ mRNA expression (Fig S2h), as well as its extracellular release (Fig. S2i). Similar enhancement of virus replication and enhanced *Ifnβ* mRNA was also observed in STING depleted mouse neuronal cell line Neuro2a (Fig. S2j-n).

We further confirmed the role of STING using H-151 which is a potent, irreversible inhibitor of STING [46]. It is known to bind covalently to the transmembrane cysteine residue of STING and blocks palmitoylation and clustering. After establishing non-toxic concentrations (Fig. S3a), the effect of 5 µM drug was checked on IFNβ activation in poly (dA: dT) treated MEFs, where a major inhibition was observed (Fig. S3b-c). In JEV infected cells H-151 treatment enhanced virus replication (Fig. S3d-g), TBK1, IRF3 activation (Fig. S3f, h-i) and downstream type-I IFN production (Fig. S3j-k). Similar increase in virus replication and IFN expression was seen in H-151 treated Neuro2a cells (Fig. S3l-p). Collectively, these data highlight that the antiviral function of STING in JEV infection is independent of IFN production.

We next investigated the impact of STING agonist diABZI on virus replication. diABZI is a novel, non-nucleotide-based STING agonist that activates STING and its downstream signaling. Cell cytotoxicity of the drug was determined using MTT assay (Fig. S4a) and a concentration of 2 µM was established as non-toxic. diABZI treatment resulted in significant activation of STING (Fig S4c) and reduction in virus replication in both MEFs and BMDMs (Fig S4a-j), underscoring the antiviral role of STING in JEV replication.

### TBK1 plays an indispensable role in mediating type-I IFN response during JEV infection

The C-terminal tail (CTT) of STING recruits TBK1, which phosphorylates S365 on STING. This then serves as a binding site for the transcription factor IRF3, which is subsequently phosphorylated and activated by TBK1, leading to transcription of IFNs and ISGs [10]. However, in JEV infection we observed enhanced TBK1 phosphorylation upon STING depletion and pharmacological inhibition (Fig. 3e-f, S2d, f, S3d, f), highlighting that TBK1 activation occurs independently of STING. We next examined the role of TBK1 in mediating JEV induced IFN response, and depleted TBK1 using siRNA mediated knockdown (Fig. 4a). TBK1 depletion enhanced virus replication (Fig. 4b-d, f), and suppressed STING phosphorylation (Fig 4c, e). Transcriptional activation of *Irf7* and *Ifnβ* was also significantly reduced (Fig. 4g-h), as well as IFNβ secretion (Fig. 4i), signifying a crucial antiviral role of TBK1 in JEV infection, likely to be mediated through type-I IFN response. To confirm the role of TBK1 we used BX-795 which is a potent and specific inhibitor of TBK1/IKKε and blocks TBK1 mediated activation of IRF3 and type-I IFN production [47]. A concentration of 250 nM was established as non-toxic and used for further experiments (Fig S5). BX-795 treatment also enhanced virus replication (Fig 4j-l,n), reduced STING phosphorylation (Fig 4k,m), and activation of IRF7 and IFNβ (Fig 4o-q), highlighting an essential role of TBK1 in mediating JEV-induced innate immune response.

**Figure 4:**
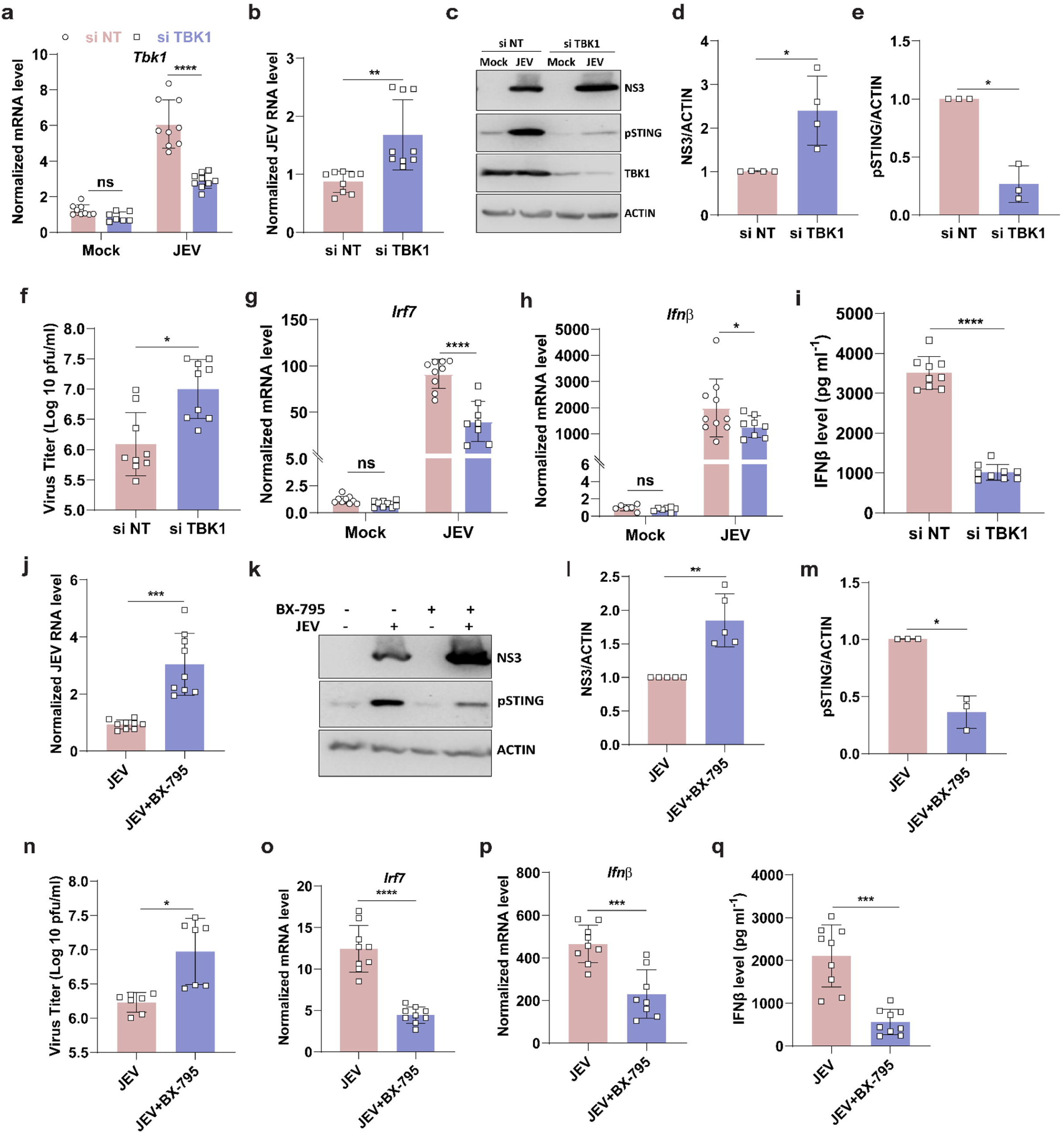
TBK1 plays an indispensable role in mediating type-I IFN response during JEV infection. NT/TBK1 siRNA (30 nM, 48 h) treated MEFs were mock/JEV infected at 2 MOI for 24 h. Total RNA was isolated to quantitate *Tbk1* mRNA expression (a) and JEV-RNA levels (b) by qRT-PCR. (c) Protein lysates were immunoblotted with JEV-NS3, pSTING, TBK1 and ACTIN (loading control) antibodies. Bar graph shows the normalized JEV-NS3(d) and pSTING (e) protein expression (JEV/mock). (f) Culture supernatant was collected to determine extracellular virus titer by plaque assay. Transcript levels of *Irf7* (g) and *Ifnβ* (h) were quantitated by qRT-PCR and normalized to si NT Mock. (i) Culture supernatant collected from mock/JEV-infected MEFs was used to perform ELISA for IFNβ. MEFs were mock/JEV infected at 2 MOI followed by treatment with BX-795 for 24 h (250 nM) (j) Cells were harvested at 24 hpi and total RNA was isolated to quantitate JEV RNA levels by qRT-PCR. (k) Protein lysates were immunoblotted with JEV-NS3, pSTING and ACTIN (loading control) antibodies. Bar graph shows the normalized JEV-NS3 (l) and pSTING (m) protein expression (JEV/mock). (n) Culture supernatant was collected to determine extracellular virus titer by plaque assay. Transcript levels of *Irf7* (o) and *Ifnβ* (p) were quantitated by qRT-PCR. (q) Culture supernatant collected from mock/JEV-infected MEFs was used to perform ELISA for IFNβ. Data presented is mean ± SD of values obtained from 3 independent experiments. Student t-test was used to calculate P values. *P < 0.05, **P < 0.01, ****P < 0.0001. ns, non-significant.

### STING mediated antiviral response is independent of autophagy

STING-induced autophagy has been reported to restrict virus replication independent of TBK1 and IFN signaling [48]. Studies from our lab have also shown that autophagy induction negatively regulates JEV replication [41, 49]. To examine if the observed antiviral role of STING was autophagy dependent, we used WT and *atg5-/-* MEFs [50]. These cells were treated with siNT/STING for 48 h followed by infection with JEV. As previously reported, autophagy deficient cells supported increased virus replication (Fig 5a, d). Additionally, in both WT and *atg5-/-* MEFs, STING depletion resulted in enhanced JEV RNA levels (Fig. 5a), viral protein (Fig. 5b, c) and titers (Fig. 5d), suggesting that STING mediated virus restriction is independent of functional autophagy machinery. Similar to our results in WT cells, we saw enhanced mRNA expression (Fig. 5e) and extracellular release (Fig. 5f) of IFNβ in STING depleted *atg5-/-* cells. We validated this finding using the STING inhibitor H-151, and the drug was able to significantly enhance virus infection and IFN response in both WT and *atg5-/-* MEFs (Fig. 5g-l). These results suggest that the antiviral role of STING is not mediated through autophagy.

**Figure 5:**
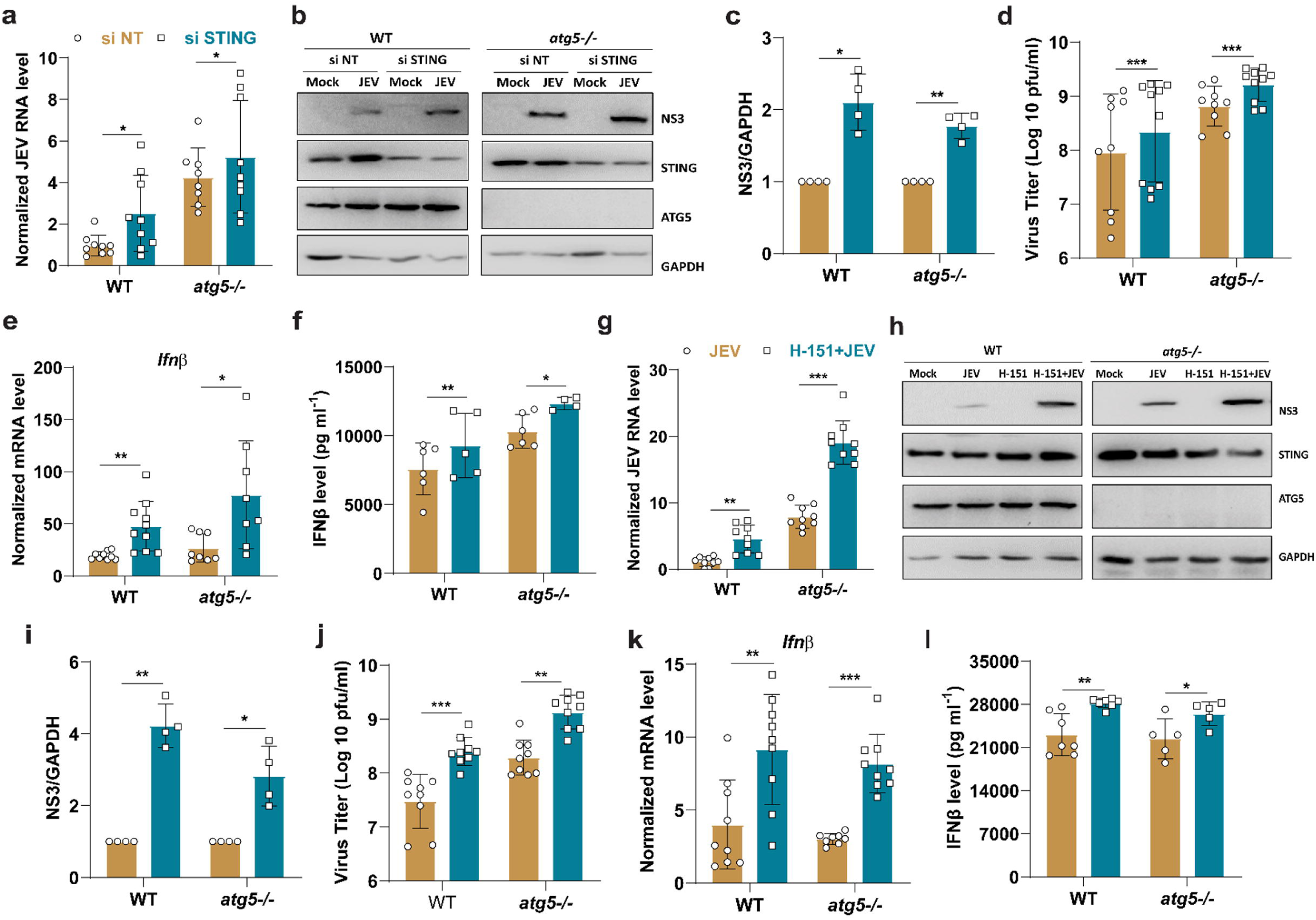
STING mediated anti-viral response is independent of autophagy. NT/STING siRNA (30 nM, 48 h) treated WT and *atg5-/-* MEFs were mock/JEV infected at 2 MOI for 24 h. Total RNA was isolated to quantitate JEV-RNA levels (a) and *Ifnβ* transcript levels (e). (b) Protein lysates were immunoblotted with JEV-NS3, STING, ATG5 and GAPDH (loading control) antibodies. (c) Bar graph shows the normalized JEV-NS3 protein expression ratio (JEV/mock) for three independent experiments. (d) Culture supernatant was collected to quantify virus titre by plaque assay and IFNβ levels (f) through ELISA. WT and *atg5-/-* cells were pretreated with DMSO or 5 µM of H-151 for 12h followed by mock/JEV infection at 2 MOI for 24 h. Total RNA was isolated to quantitate JEV-RNA levels (g) and *Ifnβ* transcript levels (k) by qRT-PCR. (h) Protein lysates were immunoblotted with JEV-NS3, STING, ATG5 and GAPDH (loading control) antibodies. (i) Bar graph shows the normalized JEV-NS3 protein expression ratio (JEV/mock) for three independent experiments. (j) Culture supernatant was collected to quantify virus titre by plaque assay and IFNβ levels (l) through ELISA. Data presented is mean ± SD of values obtained from 3 independent experiments. Student t-test was used to calculate P values. *P < 0.05, **P < 0.01, ****P < 0.0001. ns, non-significant.

### STING regulates JEV-induced non-canonical inflammasome activation and release of inflammatory cytokines

To determine the global impact of STING in JEV-infected cells, we performed an RNA-sequencing analysis of biological triplicates of mock-infected control and JEV-infected WT and STING^gt/gt^ BMDMs. JEV infection was confirmed by qRT-PCR (Fig. 6a), and plaque assay (Fig. 6b). Consistent with our earlier results with STING^gt/gt^ MEFs, BMDMs show enhanced virus infection.

**Figure 6:**
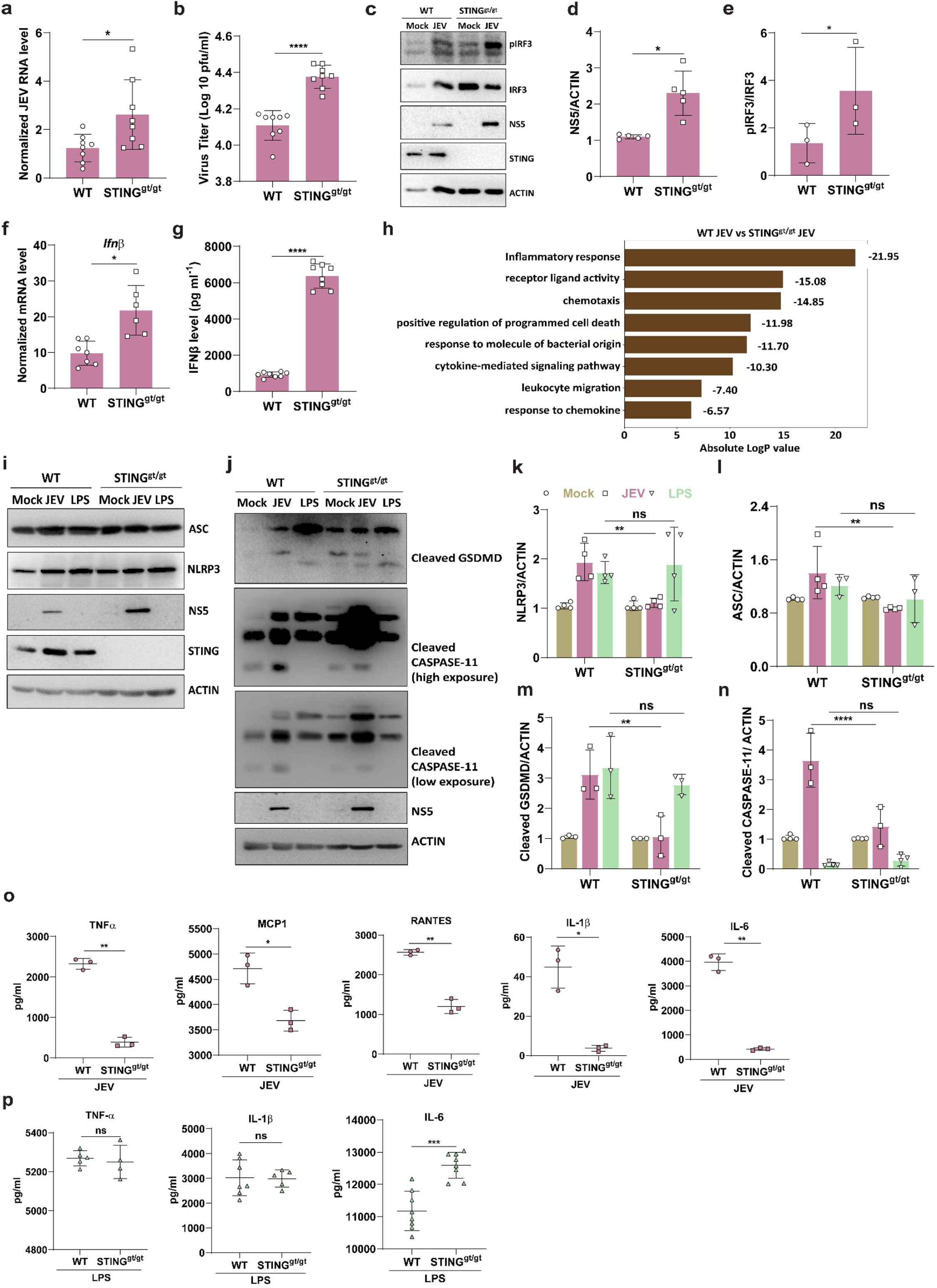
STING regulates JEV-induced non-canonical inflammasome activation and release of inflammatory cytokines. Biological triplicates of WT and STING^gt/gt^ BMDMs were mock/JEV infected (5 MOI, 24 h) (a) Total RNA was isolated to quantitate JEV-RNA levels. (b) Culture supernatant was collected to determine extracellular virus titer by plaque assay. (c) Protein lysates were immunoblotted with pIRF3, IRF3, JEV-NS5, STING and ACTIN (loading control) antibodies. Bar graph shows the normalized NS5/ACTIN (d) and pIRF3/IRF3 (e) protein expression ratio (JEV/mock) for three independent experiments. Total RNA was isolated to quantitate the transcript levels of *Ifnβ* (f). (g) Culture supernatant collected from mock/JEV-infected BMDMs was used to perform ELISA for IFNβ. (h) Pathway enrichment analysis of the downregulated DEGs in JEV-infected STING^gt/gt^ BMDMs compared to WT BMDMs. BMDMs isolated from WT and STING^gt/gt^ C57BL/6 mice were either mock/JEV infected at 5 MOI for 24 h or treated with LPS (1 mg/ml) for 4 h followed by nigericin (1 µM) for 1 h. (i,j) Protein lysates were immunoblotted with NLRP3, ASC, GSDMD, CASPASE-11, JEV-NS5 and ACTIN (loading control) antibodies. Bar graph shows the normalized NLRP3, ASC, cleaved CASPASE-11 and cleaved GSDMD (k,l,m,n) protein expression (JEV/mock) for three independent experiments. (o) Cytokines (TNF-α, MCP-1, RANTES, IL-1β and IL-6) were quantified from the culture supernatant using CBA (n=3). (p) WT and STING^gt/gt^ C57BL/6 BMDMs were treated with LPS (1 mg/ml) for 4 h followed by nigericin (1 µM) for 1 h. Culture supernatant was collected to determine cytokine levels of TNF-α, IL-1β and IL-6 through ELISA. Data presented is mean ± SD of values obtained from 3 independent experiments. Student t-test was used to calculate P values. *P < 0.05, **P < 0.01, ****P < 0.0001. ns, non-significant.

JEV infected STING^gt/gt^ BMDMs showed enhanced expression of key genes involved in innate and adaptive immune response to virus (Fig. S5a). Critical transcripts of innate immune signaling pathway particularly type-I IFN response (*Ifnb1, Ifna1, Ifna2, Ifnab, Gm13276*) and T cell activation (*Cxcr4, Egr1, Mdk, Rora*) were found to be highly upregulated in JEV-infected STING^gt/gt^ BMDMs (Fig. S5b). We also validated STING mediated regulation of downstream adaptors and observed enhanced IRF3 phosphorylation (Fig. 6c, e), viral NS5 protein level (Fig. 6c-d) and *IFNβ* mRNA expression (Fig. 6f) and release (Fig. 6g). The transcriptomics data also showed upregulation of several inflammatory cytokines and chemokines in WT BMDMs (Fig. S5c). However, JEV-infected STING^gt/gt^ BMDMs exhibited significant downmodulation of genes associated with inflammatory responses, cytokine mediated signaling pathway and programmed cell death (Fig. 6h). Critical transcripts of genes involved in acute inflammatory response (*Il1a, Il1b, Il6, Ccl5, Cxcl1*) and cytokine production, programmed cell death (*Lcn2, Sod2, Map2k6*) and chemotaxis (*Cxcl3, Ccl5, Ccl7, Cxcl2*) were found to be suppressed in JEV-infected STING^gt/gt^ BMDMs (Fig. S5d).

We next attempted to functionally validate the activation of inflammasome and cell death markers in WT and STING^gt/gt^ BMDMs with LPS as a positive control. Key proteins involved in inflammasome activation and pyroptotic cell death including NLRP3, ASC, cleaved CASPASE-11 and cleaved Gasdermin D (GSDMD) (Fig. 6i-n,) were highly upregulated in both JEV infected and LPS stimulated WT BMDMs. Strikingly, in JEV infected STING^gt/gt^ BMDMs protein levels of all these key markers was significantly less compared to infected WT BMDMs. Additionally, the release of pro-inflammatory cytokines and chemokines like TNF-α, MCP-1, RANTES, IL1-β and IL-6 (Fig. 6o) was also significantly impaired in STING^gt/gt^ BMDMs compared to WT cells. Overall, these data demonstrate that STING^gt/gt^ BMDMs are defective in their ability to produce and secrete pro-inflammatory cytokines in response to JEV infection. However, we found no significant difference in the ability of LPS induced STING^gt/gt^ BMDMs to activate inflammasome and cell death pathway proteins as well as release of pro-inflammatory cytokines (Fig. 6i, j, p), suggesting that these cells are not intrinsically impaired in generating inflammatory cytokines in response to other external stimuli.

### STING recruits NLRP3 to the virus replication complex

There is evidence in literature that STING interacts with NLRP3, promotes its localization in the ER and triggers inflammasome activation [13, 15]. Since STING was found to localize with virus replication membranes, we tested the localization of STING and NLRP3 in JEV infected cells. We observed that in uninfected cells, NLRP3 was diffusely distributed in the cytosol, however upon JEV infection it formed distinct puncta and colocalized with pSTING on the virus replication complex (Fig. 7a). We next attempted to examine if this NLRP3 recruitment on virus replication complex is STING dependent. For that, we generated STING KO MEFs using CRISPR-Cas9 system (Fig. 7b). While, WT cells showed NLRP3 recruitment on the RC, the STING deficient cells showed diffused distribution of NLRP3 in the cytosol (Fig. 7c). These data clearly indicate that STING is essential for localization of NLRP3 on the virus replication complex and serves as a hub for recruitment of immune effectors and inflammasome activation.

**Figure 7:**
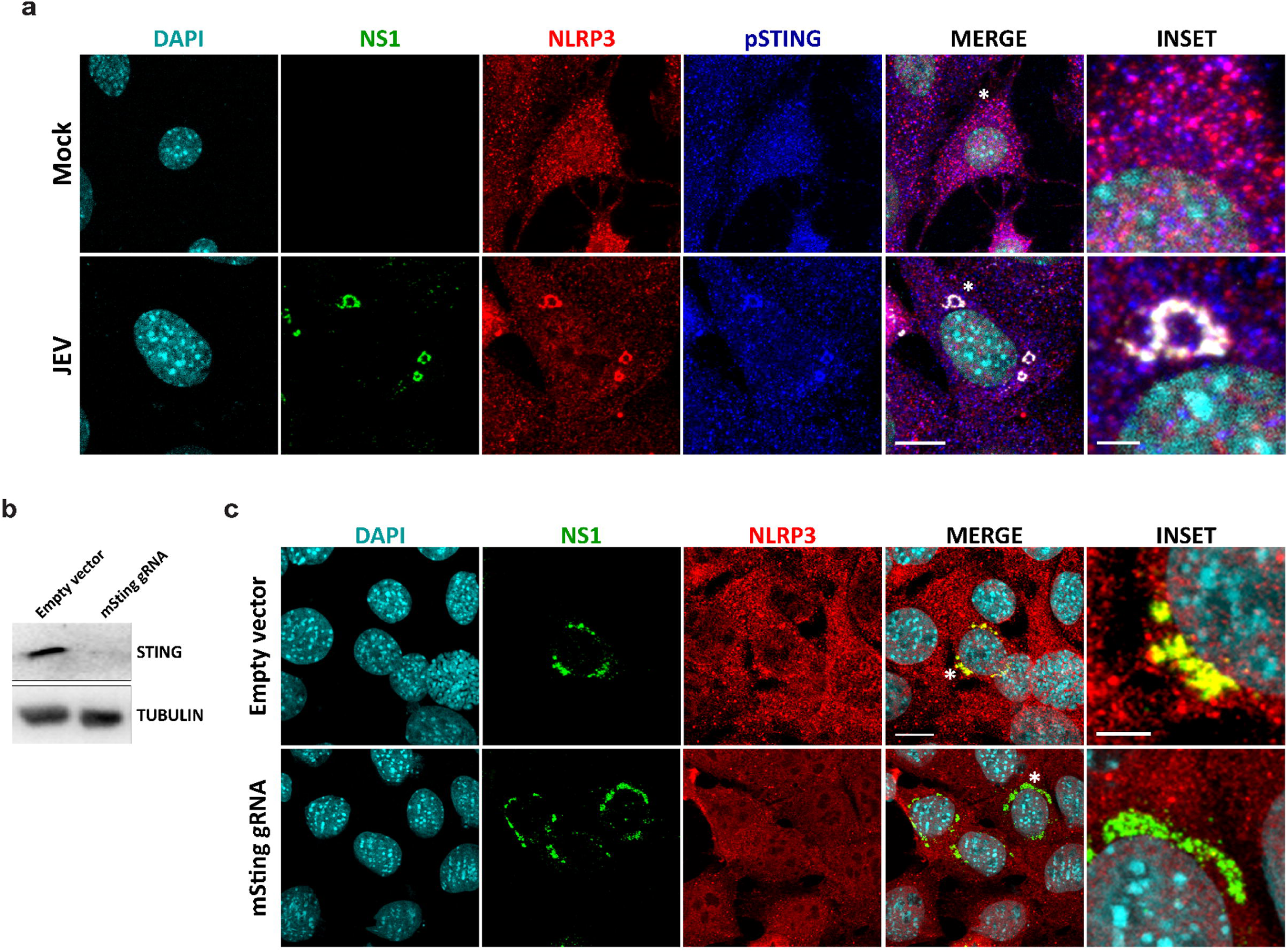
STING recruits NLRP3 to the virus replication complex. MEFs were mock/JEV infected with JEV (5 MOI, 24h). Cells were stained (a) with JEV-NS1 (green), NLRP3 (red) and pSTING (blue) antibodies (c) or with JEV-NS1 (green) and NLRP3 (red) antibodies. Nucleus was stained with DAPI. Images were acquired on a confocal microscope using a 63x objective. Right most panel shows the magnified area marked by asterisk (*). Scale bar 10µm. Images are representative of two independent experiments. (b) Protein lysates were immunoblotted with STING and TUBULIN (loading control) antibodies.

### STING proton channel activity mediates JEV-induced NLRP3 inflammasome activation

Since canonical STING-TBK1-IRF3 axis mediates NF-κβ/IKK activation thereby inducing production of pro-inflammatory cytokines and subsequent innate inflammation, we assessed the phosphorylation status of NF-κβ and IKKε. Enhanced phosphorylation of NF-κβ/IKKε was seen in STING^gt/gt^ cells, suggesting NF-κβ/IKK axis was dispensable for STING mediated induction of inflammatory cytokines during JEV infection (Fig. 8a-c).

**Figure 8:**
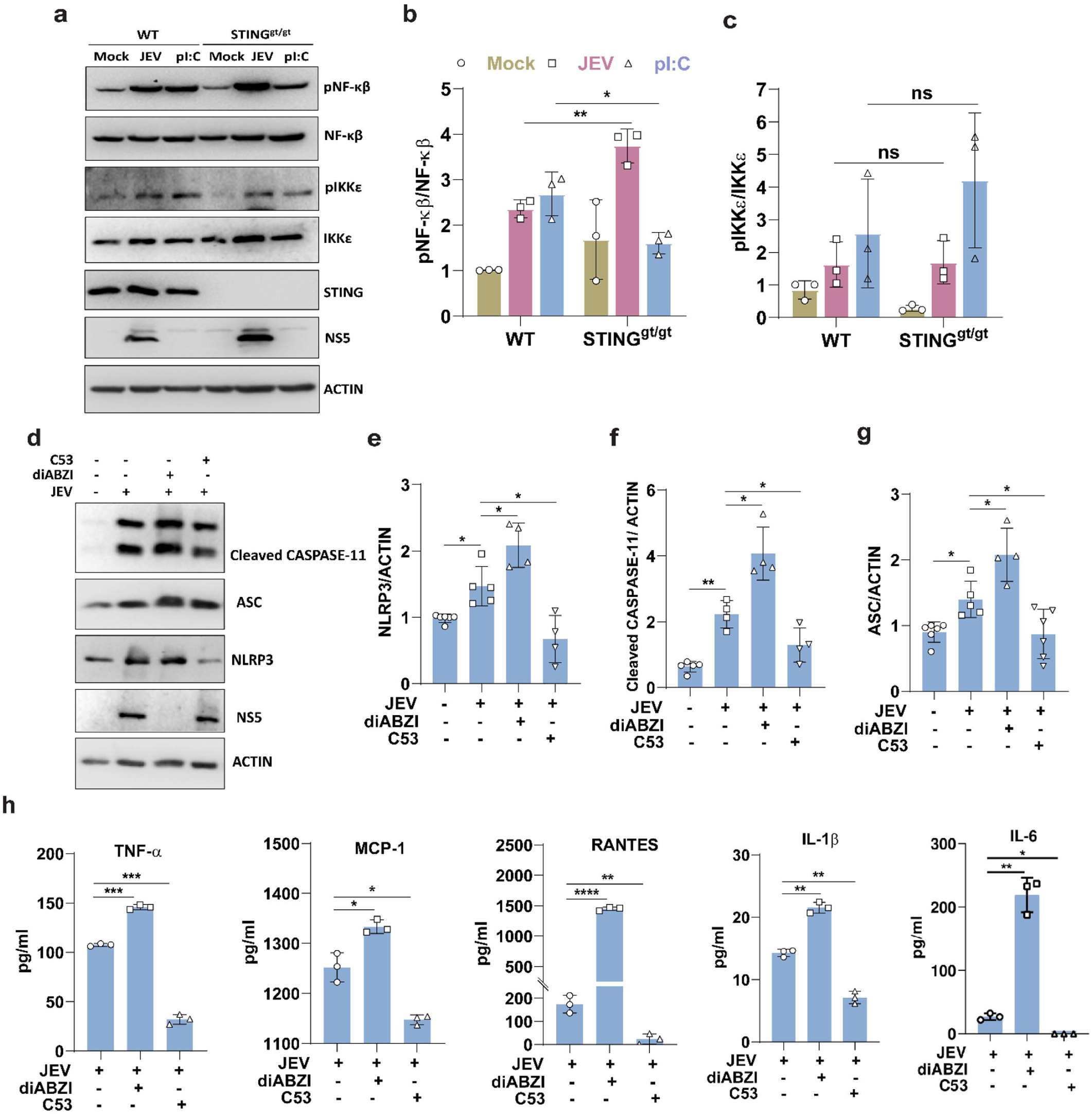
STING proton channel activity mediates JEV-induced NLRP3 inflammasome activation. WT and STING^gt/gt^ MEFs were either mock/JEV infected at 2 MOI for 24 h or transfected with pI:C (1 μg/ml) for 6 h. (a) Protein lysates were prepared and immunoblotted with pNF-κβ, NF-κβ, pIKKε, IKKε, and ACTIN (loading control) antibodies. (b,c) Bar graph shows the normalized pNF-κβ/NF-κβ and pIKKε/IKKε protein expression (JEV/mock) for three independent experiments. BMDMs were either pre-treated with C53 (10 µM) for 2 h, then mock/JEV infected for 24 h, or co-treated with diABZI (2 µM) along with mock/JEV infection for 24 h.(d) Protein lysates were immunoblotted with CASPASE-11, ASC, NLRP3, JEV-NS5 and ACTIN (loading control) antibodies. (e,f,g) Bar graph shows the normalized NLRP3, cleaved CASPASE-11 and ASC protein expression compared to Mock. (h) Cytokines (TNF-α, MCP-1, RANTES, IL-1β and IL-6) were quantified from the culture supernatant using CBA (n=3).

Recent findings have reported that upon activation, STING induces proton leakage at the Golgi by forming a channel at the interface of its transmembrane domain homodimer. This then regulates non canonical autophagy induction and Golgi transit as well as secretion of inflammatory cytokines [51, 52]. To test if the proton channel activity of STING regulates JEV-induced inflammasome activation, we used a small-molecule agonist compound 53 (C53) which binds to the cryptic site in the STING transmembrane domain and inhibits its channel activity. We first determined the non-toxic concentration of C53 in BMDMs which was found to be 10 µM and used for further experiments (Fig. S7a). Treatment of BMDMs with the STING agonist diABZI post JEV infection resulted in enhanced NLRP3 inflammasome activation and its downstream mediators (ASC and cleaved CASPASE-11), which was all blocked upon C53 treatment (Fig. 8d-g). Alongside inflammatory cytokine levels were enhanced upon STING activation with diABZI treatment and significantly impaired upon proton channel inhibition using C53 treatment (Fig. 8h). Additionally, diABZI and C53 did not affect pyroptotic cell death markers (Fig. S7b-c) and cytokine release (Fig. S7d) in LPS and nigericin stimulated cells. Collectively, these results support that in JEV infection, NLRP3 inflammasome activation occurs through STING proton channel-dependent mechanisms.

### GSDMD is a critical host factor for defense against JEV infection

Our results clearly suggested that STING plays a crucial antiviral role by regulating the activation of NLRP3 inflammasome and pyroptotic cell death pathway markers. GSDMD, the primary effector of pyroptosis, facilitates the release of pro-inflammatory molecules into the extracellular space through its pore-forming activity [53]. Therefore, we checked the role of pyroptotic cell death pathway proteins CASPASE-11 and GSDMD during JEV infection. MEFs were treated with siNT/CASPASE-11 for 48 h (Fig. S8a), and this resulted in significantly enhanced JEV RNA replication (Fig. S8b-e). Similarly, in the mouse microglial cell line N9, CASPASE-11 knockdown resulted in enhanced virus replication (Fig. S8f-j). We next depleted GSDMD (Fig. 9a) and observed a marked enhancement of JEV replication (Fig 9b-e). Upon GSDMD depletion the transcript levels *of Ifnβ and Il1β* were greatly reduced (Fig. 9f). Similar observations were made in microglial N9 cells, where GSDMD depletion was proviral (Fig S9a-e), but decreased the transcript levels of chemokines and cytokines including *Cxcl10* and *Il1β* (Fig S9f), and IL-1β production (Fig. S9l). Interestingly, GSDMD also localized with virus replication complexes and NLRP3 (Fig. 9g). These results clearly highlight that the virus replication membranes are the focal-point of the STING mediated inflammasome assembly which recruits effectors such as GSDMD and restricts virus replication.

**Figure 9:**
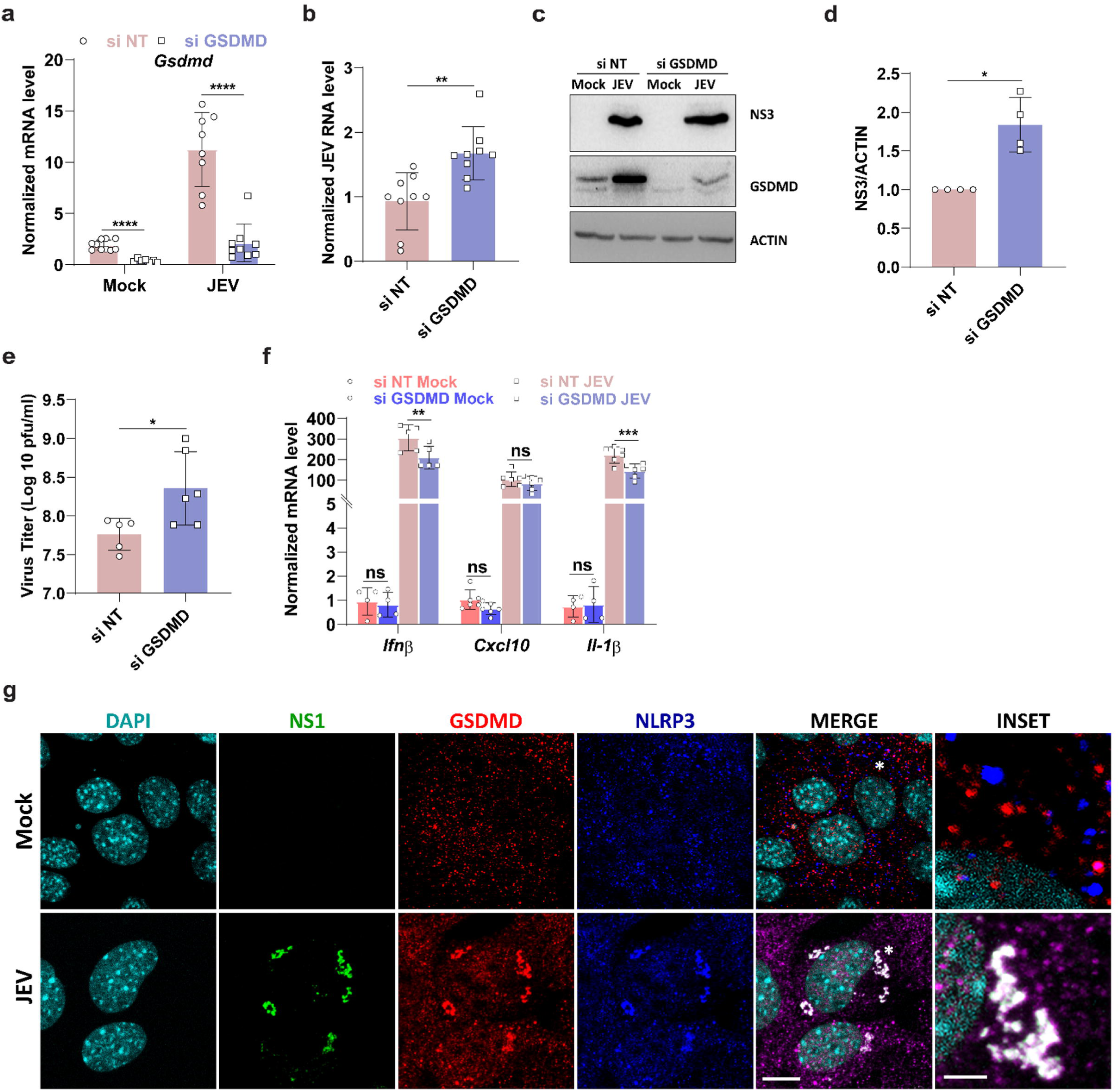
GSDMD is a critical host factor for defense against JEV infection. NT/GSDMD siRNA (30 nM, 48 h) treated MEFs were mock/JEV infected at 2 MOI for 24 h. Total RNA was isolated to quantitate *Gsdmd* mRNA expression (a) and JEV-RNA levels (b) by qRT-PCR. (c) Protein lysates were immunoblotted with JEV-NS3, GSDMD and ACTIN (loading control) antibodies. (d) Bar graph shows the normalized JEV-NS3 protein expression (JEV/mock). (e) Culture supernatant was collected to determine extracellular virus titer by plaque assay. (f) Transcript levels of *IFNβ, Cxcl10 and Il-1β* were determined by qRT-PCR. MEFs were mock/JEV infected with JEV (5 MOI, 24h). Cells were stained with (g) JEV-NS1 (green) and GSDMD (red) and NLRP3 (blue) antibodies. Nucleus was stained with DAPI. Images were acquired on a confocal microscope using a 63x objective. Right most panel shows the magnified area marked by asterisk (*). Scale bar 10µm. Images are representative of two independent experiments.

### Goldenticket (STING^gt/gt^) mice exhibit enhanced susceptibility to JEV infection and reduced inflammation

Finally, we tested JE disease in the Goldenticket (STING^gt/gt^) mouse model. We used mouse-adapted JEV-S3 isolate, which induces characteristic encephalitis symptoms, including weight loss, piloerection, body stiffening, and hind limb paralysis by 5–6 days post-infection (dpi), followed by death within 2–3 days of symptom onset [54]. Notably, Goldenticket (STING^gt/gt^) mice showed earlier symptoms and begin to die at 6 days post viral infection with significantly reduced survival (Fig. 10a, b) and higher viremia in the brain (Fig. 10c, d, g) compared to WT mice, suggesting STING^gt/gt^ mice are more susceptible to virus infection. We also observed significantly enhanced IFNβ production in STING^gt/gt^ mice brain as a consequence of higher viremia (Fig. 10e). Since JEV infection induces neuroinflammation, we checked the levels of key inflammatory mediators. Strikingly, we observed significantly reduced NLRP3 protein in STING^gt/gt^ mice brain lysates (Fig. 10f, h). Similarly, the cleaved form of both GSDMD and CASPASE-11 was also greatly reduced in the null mice (Fig. 10i, j, k). Consistently, JEV infected STING^gt/gt^ mice showed lower levels of pro-inflammatory cytokines including TNF-α, MCP-1, RANTES, IL-1β and IL-6 (Fig. 10l) suggesting that STING deficiency results in the attenuation of inflammation and the promotion of JEV replication in mice brain. This observation was also recapitulated in primary mixed glial cells, which are a key source of pro-inflammatory cytokines in JEV-induced CNS inflammation. The transcript levels of *Il6*, *Tnf-α, Mcp-1 and Il-1β* were reduced in JEV infected STING^gt/gt^ primary mixed glial cells. (Fig. S10a-d). JEV induced NLRP3 inflammasome activation was also greatly reduced in STING^gt/gt^ cells (Fig. S10e). Collectively, these results highlight that STING is essential for host defense against JEV infection by facilitating inflammatory responses.

**Figure 10:**
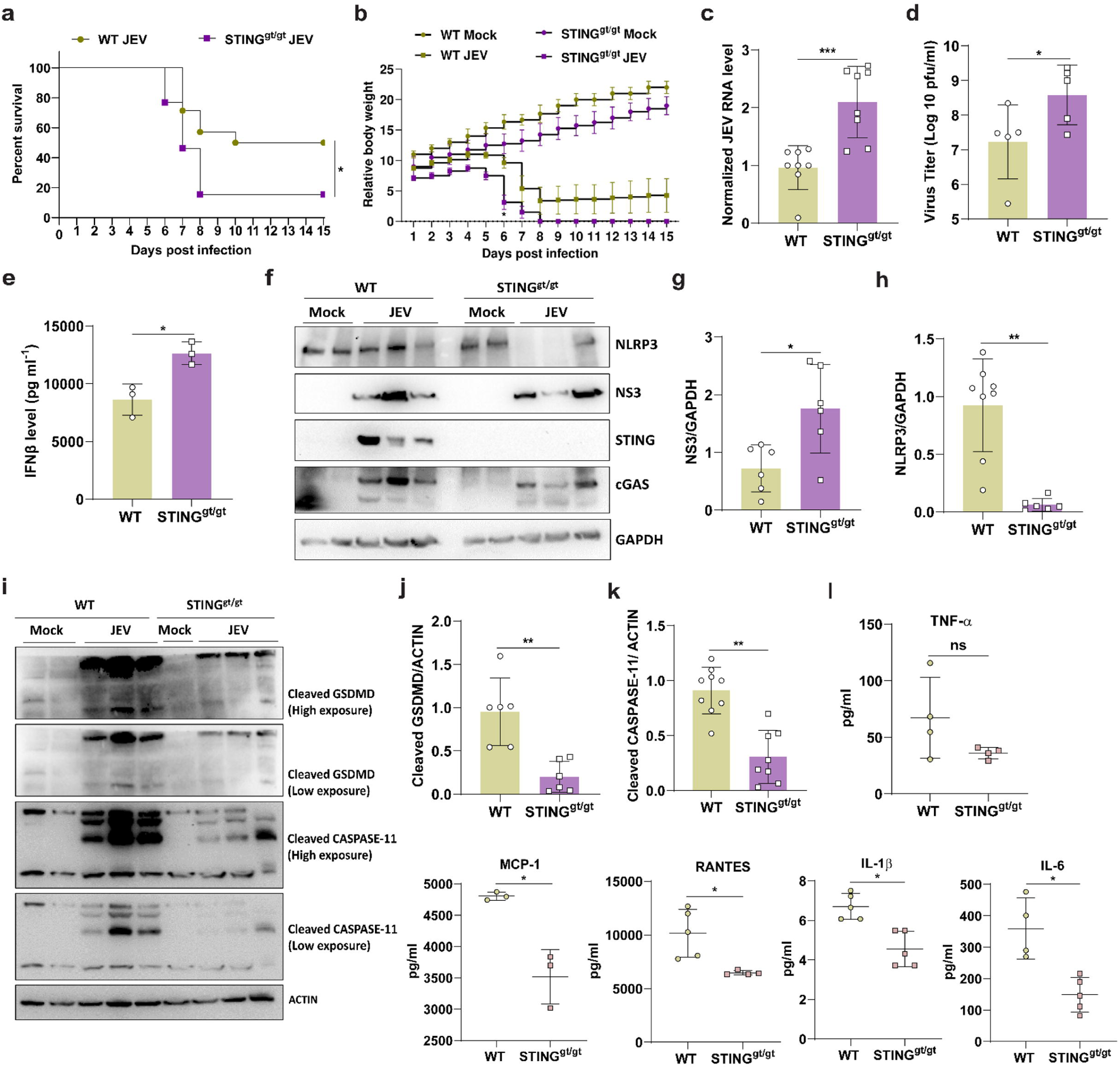
Goldenticket (STING^gt/gt^) mice exhibit enhanced susceptibility to JEV infection and suppressed pyroptotic cell death. Three-week-old C57BL/6 and STING^gt/gt^ mice were mock/JEV-S3 (10^7^ pfu) infected through an i.p. injection. All mice were monitored for the appearance of encephalitis symptoms until death. (a) Survival curve of mock (n=3)/JEV (n=14) in each group. Log-rank (Mantel-Cox) test was used to determine the statistical significance comparing WT JEV and STING^gt/gt^ JEV mice group (b) Graph representing the change in body weight of JEV infected WT and STING^gt/gt^ mice group normalized to mock-infected mice group, compared by unpaired Student t-test. Mock/JEV infected mice were sacrificed. Brain tissues were divided into three equal parts and homogenized. Total RNA was isolated from one part. JEV RNA levels (c) were determined by qRT-PCR. (d) Culture supernatant was collected to quantify virus titre by plaque assay (e) and IFNβ levels (g) through CBA. (f) Protein lysate was prepared from another part and western blot analysis was performed with NLRP3, JEV-NS3, STING, cGAS and ACTIN (loading control) antibodies. Bar graph shows the normalized JEV-NS3 (g) and NLRP3 (h) protein expression ratio (JEV/mock). (i) Protein lysates were immunoblotted with GSDMD, CASPASE-11 and ACTIN (loading control) antibodies. (j,k) Bar graph shows the normalized cleaved CASPASE-11 and cleaved GSDMD protein expression ratio (JEV/mock) (l) 30µg protein from each sample was used for the quantitation of cytokine levels using CBA. Data were analyzed with LEGENDplexTM Multiplex assay software. Data presented is mean ± SD of values. Student t-test was used to calculate P values. *P < 0.05, **P < 0.01, ****P < 0.0001. ns, non-significant *P < 0.05, **P < 0.01, ****P < 0.0001. ns, non-significant.

**Figure 11:**
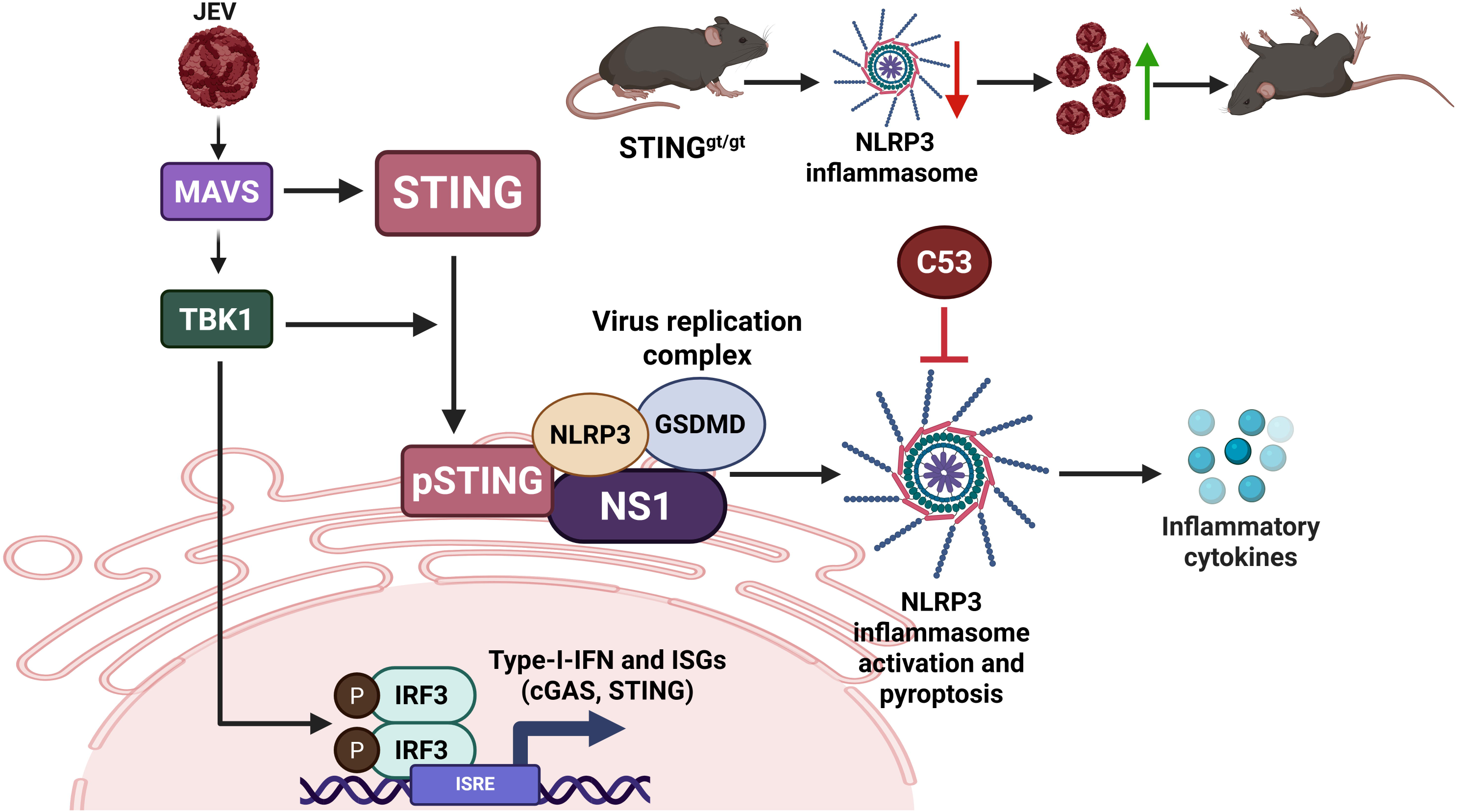
Schematic model summarising the activation and role of STING during JEV infection. STING gets activated in virus-infected cells downstream of MAVS and IFN signaling, and is recruited to the virus replication complex, which acts as a hub for inflammasome activation, release of proinflammatory cytokines, and induction of pyroptotic cell death markers. The proton channel activity of STING is crucial for inflammatory response. STING is neuroprotective, as the Golden ticket null-animal model displays increased viremia, lower inflammation and reduced survival. Created with Biorender.com

## Discussion

The host innate immune system provides the first line of defense against microbial infection. Macrophages, dendritic cells as well as non-professional cells like epithelial, endothelial and fibroblast cells play important roles in detecting pathogen with the help of PRRs [34, 55]. cGAS is a PRR which senses cytosolic DNA and catalyses the synthesis of 2ʹ3ʹ-cyclic GMP–AMP (cGAMP), a secondary messenger that directly binds and activates STING, initiating a signaling cascade through TBK1 and IRF3 that culminates in type I IFN production [10, 24, 56, 57]. In contrast, RNA viruses are primarily detected by the membrane-bound TLRs localized to the ER and endosomes (e.g., TLR3, TLR7/8), as well as by cytosolic DExD/H-box RNA helicases, including RIG-I and melanoma differentiation-associated gene 5 (MDA5, also known as IFIH1) which signal through MAVS to activate the same TBK1–IRF3 axis [58, 59]. While the involvement of cGAS-STING axis in sensing DNA viruses is well understood, recent evidences have emerged highlighting their role during RNA virus infection [29, 60]. Although direct activation of cGAS by viral RNA is unlikely, as RNA binding does not induce cGAMP production [61], indirect activation of the cGAS/STING pathway through non-canonical mechanisms has been observed [62, 63]. RNA viruses from diverse families—such as Alphaviruses, Flaviviruses, Orthomyxoviruses, and human coronaviruses (hCoVs) have been shown to interact with and modulate STING signaling [33, 34, 37, 64–66].

Proteomic study from our lab on JEV infected fibroblasts demonstrated robust activation of cGAS-STING signaling pathway and an antiviral role of cGAS [8]. Here, we investigated the role of its downstream adaptor STING. JEV infection activated STING, independent of cGAMP production. Activated STING localised to the virus replication membranes and served as a hub for NLPR3 recruitment and inflammasome activation. This manifested in an antiviral and neuroprotective effect, evidenced in the Goldenticket (STINGgt/gt) mouse model.

Various host factors [33] and viral proteins [66] have been shown to regulate the expression of cGAS-STING at both transcriptional and posttranslational levels; however, the regulatory mechanisms remain incompletely characterized. Notably, viral components such as open reading frames and non-structural proteins from SARS-CoV-2, the Vif protein from HIV-1, and the papain-like protease (PLP) from HCoV-NL63 have been implicated in regulating cGAS– STING activity [65, 67]. As reported by Ma et al, type-I IFN stimulation in BMDMs induces *cgas* & *Sting* mRNA as well as protein expression, identifying cGAS-STING as ISGs [68, 69] which in turn mediate a positive feedback loop for type-I IFN production. Studies have also demonstrated that STING is a late-induced gene downstream of RIG-I-MAVS activation, driven by autocrine and paracrine signaling through the synergistic action of tumor necrosis factor-alpha (TNF-α) and IFNs [70, 71]. As already reported, RIG-I-MAVS pathway gets activated during JEV infection and plays an important role in mediating JEV induced immune and inflammatory responses [7]. We also observe that MAVS-IFN signaling is critical for mounting an effective immune response and restricting JEV replication. Further, our results also implicate that cGAS and STING mRNA as well as protein expression during JEV infection is regulated by MAVS-IFN signaling pathway highlighting their role as ISGs. Taken together these results suggest that there is a crosstalk between RNA sensing RIG-I-MAVS pathway and DNA sensing cGAS-STING pathway during JEV infection.

Some reports have suggested a role of STING in the induction of IFNs and ISGs during RNA virus infection [60, 63, 64, 72, 73], however other IFN independent antiviral mechanisms including induction of autophagy [67, 74] and viral protein translation inhibition [29] have also been reported. We first examined the role of STING in establishing an antiviral state inside the cell post JEV infection. Knockdown, null mutant and inhibitor studies revealed that STING-deficient cells retained the ability to induce type-I IFN during JEV infection. However, despite their ability to activate TBK1-IRF3 to induce IFN response as well as ISGs, they were unable to restrict JEV replication suggesting STING plays an antiviral role through IFN independent mechanisms. The enhanced type-I IFN production upon STING deficiency may be a consequence of increased viremia that triggers STING-independent innate immune sensing pathways such as TLRs and RIG-I. Our findings with TBK1 depletion supported this idea that TBK1-IRF3 get activated through STING independent pathways during JEV infection and mediate antiviral innate immune response via production of IFN and ISGs.

The primordial function of STING discovered by Gui et al is autophagy induction that is independent of TBK1 activation and IFN signaling [11, 48]. Recently, STING has also been discovered as a proton channel, which drives LC3B lipidation onto single membrane vesicles mediated by recruitment of V-ATPase and WD40 domain of ATG16L1, independently of the canonical upstream autophagy machinery [51, 75]. STING-induced autophagy has been reported to restrict *M. tuberculosis* infection via lysosomal degradation of the pathogen. [76]. Lately, it was also reported that STING effectively activated non-canonical autophagy to suppress HSV-1, followed by viral degradation through the lysosomal pathway [30]. Our earlier studies have established that autophagy induction negatively regulates JEV replication [41, 45]. These studies prompted us to examine the role of autophagy machinery in regulating STING mediated antiviral response. Consistent with the previous reports, we observed that JEV replication was enhanced in autophagy deficient cells. However, both knockdown and inhibitor studies in *atg5-/-* cells demonstrated that STING mediates an autophagy-independent antiviral effect during JEV infection.

In the periphery, JEV mainly infects and replicates in various immune cells including monocytes/macrophages and dendritic cells. These cells are responsible for eliciting an effective immune response in the periphery which clears the virus in most of the cases [77]. Hence to further explore the antiviral mechanism, we performed RNA sequencing analysis of mock-infected control and JEV-infected BMDMs isolated from WT and STING^gt/gt^ C57BL/6 mice. As has been previously reported, our transcriptomic data also revealed upregulation of genes involved in innate immune response, inflammatory response, T-cell proliferation as well as cellular response to interferons upon JEV infection. Additionally, in STING^gt/gt^ BMDMs we observed further enhanced transcription of genes involved in innate and adaptive immune response to virus with major pathways involving positive regulation of peptidyl-serine phosphorylation of STAT protein, response to dsRNA, natural killer cell activation, B cell proliferation and T cell activation. These results again implicated STING is dispensable for JEV induced innate immune response.

There is also evidence in literature that JEV infection activates diverse programmed cell death pathways particularly apoptosis, pyroptosis, and necroptosis. Transcriptome profile of JEV infected macrophages revealed increased expression of cell death markers and inflammatory cytokines including Caspase-4, NLRP3, AIM-2, Caspase-1, Gasdermin D (GSDMD), IL-1β, and IL-18 [78]. STING activation also results in phosphorylation and nuclear translocation of NF-κβ/IKKε axis which induces production of inflammatory cytokines and chemokines. Non-canonically, cGAS-STING pathway has also been associated with the activation of NLRP3 inflammasome as well as production of pro-inflammatory cytokines and chemokines [14–16, 79]. Effector memory T cells have been reported to induce innate inflammation by causing DNA damage in dendritic cells, which activates a cGAS-independent, non-canonical STING– TRAF6–NF-κB pathway leading to pro-inflammatory cytokine production and autoimmunity [80]. During acute kidney injury, STING mediated ER and oxidative stress contributes to tubular cell inflammation and pyroptosis [81]. Yang et al demonstrated that STING deficiency alleviates diabetic myopathy by inhibiting NLRP3-mediated pyroptosis [82]. STING has also been shown to enhance NLRP3 inflammasome-mediated hepatocyte pyroptosis during liver injury via epigenetic upregulation through the IRF3/WDR5/DOT1L complex, contributing to liver fibrosis by enhancing inflammation and altering hepatic metabolism [83]. In our study, we observed significant downregulation of genes involved in inflammatory responses, cytokine mediated signaling pathway, IL-17 signaling pathway, NLR signaling pathway and programmed cell death in JEV infected STING^gt/gt^ BMDMs. Functional validation demonstrated reduced activation of key proteins involved in NLRP3 inflammasome and pyroptotic cell death pathway including NLRP3, ASC, cleaved CASPASE-11 as well as cleaved GSDMD in STING^gt/gt^ cells. Furthermore, the extracellular release of pro-inflammatory cytokines was also impaired in these cells indicating an essential role of STING in activating NLRP3 inflammasome. The reduced levels of chemokines and cytokines in STING^gt/gt^ BMDMs might negatively regulate the migration of leucocyte, neutrophil and granulocyte to the inflammatory sites resulting in enhanced virus replication. There is evidence in literature that the proton channel activity of STING is required for NLRP3 inflammasome activation and cytokine release [51, 52]. Our data with C53 treatment also implicated that STING’s proton channel activity mediates NLRP3 inflammasome activation and release of pro-inflammatory cytokines and chemokines during JEV infection.

STING trafficking to the membranous virus replication organelles has been reported in context to human rhinovirus (HRV) infection, where it interacts with PI4P to facilitate virus replication and transmission [84]. Flaviviruses replicate on ER-derived membranes enriched with host factors and viral non-structural proteins [38, 39, 85]. Our results establish that STING localizes to the viral replication complex and facilitates the recruitment of NLRP3. We also observed GSDMD localization. Our earlier study has established that the ISG: Guanylate binding protein (GBP1) is also upregulated and localized to replication complexes. [86]. The concentration of these effectors could be a potential mechanism to disintegrate the replication complexes, and warrants further investigation. Collectively, these findings highlight a critical role for STING at the viral replication sites, which serve as hub for the recruitment of host defense factors.

The inflammasome and pyroptotic cell death pathway proteins serve as critical components of the innate immune system, essential for mounting effective antiviral responses against several viruses including SARS-CoV [87], EMCV and VSV [88] and HSV-1 [13]. In our study, CASPASE-11 and GSDMD were observed to be antiviral by regulating the secretion of inflammatory cytokines and chemokines.

In conclusion, our findings demonstrate that STING activation during JEV infection is regulated by the MAVS-IFN signaling axis. It plays an IFN and autophagy independent antiviral role that is largely driven by inflammation. We establish that activated STING localizes to the viral replication complex, where it facilitates inflammasome activation, promotes the release of proinflammatory cytokines and chemokines, and triggers pyroptotic cell death. STING is thus a critical neuroprotective host factor that modulates the inflammatory response during JEV infection.

## Materials and methods

### Ethics statement

Animal handling and experiments were performed as per the guidelines of the Committee for the Purpose of Control and Supervision of Experiments on Animals (CPCSEA), Government of India. Animals were maintained under a controlled 14-hour light/10-hour dark cycle at a temperature range of 19–26 °C and relative humidity of approximately 30–70%. Food and water were provided ad libitum. (RCB/IAEC/2022/127, RCB/IAEC/2024/203, and CCMB/IAEC/48/2023).

### Cell lines and virus

WT and *atg5-/-* Mouse embryonic fibroblasts (MEFs) were obtained through the RIKEN Bio-Resource Cell Bank (RCB2710 and RCB2711). C6/36 (insect), Neuro2a (mouse neuroblastoma), and Vero cell lines were obtained from the cell repository at the National Centre for Cell Sciences Pune, India. N9 (mouse microglia) cell line was a gift from Prof. Anirban Basu (NBRC, India). All cell lines were negative for mycoplasma.

MEFs and Neuro2a cells were cultured in Dulbecco’s modified Eagle’s medium (DMEM) supplemented with 10% fetal bovine serum (FBS), Vero cells were cultured in Eagle’s minimal essential medium (MEM), Leibovitz’s (L-15) medium with 10% FBS was used to culture C6/36 cells and RPMI for N9 cells. Furthermore, all media were additionally supplemented with 100 μg/ml penicillin-streptomycin, and 2 mM L-glutamine.

JEV isolate Vellore P20778 strain (GenBank accession no. AF080251) generated in C6/36 cell line was used in this study. UV-inactivated JEV was generated by exposing the virus to UV radiation (1600×100 µJ/cm²) for 20 minutes on ice. For animal experiments, the mouse-adapted JEV-S3 strain was used [54]. Virus titer was determined using Vero cells by performing plaque assays.

All reagents, antibodies and siRNA used in the study are listed in supplementary table S1.

### Generation of STING KO MEFs

STING KO MEFs were generated using a CRISPR-Cas9 editing system using pSpCas9(BB)-2A-Puro (PX459) V2.0 (Addgene, #62988) and Sting1_sgRNA1_pSpCas9 BB-2A-Puro (PX459) v2.0 (Addgene, #62988) vectors as described by Ran et al, (2013) [89]. The guide RNA (gRNA) target gene sequences viz. mouse Sting 5’-AGCGGTGACCTCTGGGCCGT-3’ cloned into PX459 vector, was procured from Genescript. The final constructs were named as PX459-gRNA-Sting1. MEFs were transfected with 2 µg of plasmids using LipofectamineTM 3000 (InvitrogenTM) following the manufacturer’s instructions. The empty vector PX459 was used as a control. At 48 hours post-transfection, knock out cells were selected by treating with puromycin (2 µg/ml) for 8 days. The knockout was validated by western blotting.

### Primary cell culture BMDMs

C57BL/6/AG129/STING^gt/gt^ mice (6-9 weeks old) were used to isolate BMDMs. Mice were euthanized in CO_2_ followed by dissection to isolate femur and tibia which were then washed with PBS and flushed with RPMI media supplemented with L929 conditioned media to collect bone marrow cells. Following RBC lysis, the cells were grown in complete RPMI media additionally supplemented with L929-conditioned media for 7 days with media changes on day 3 and 5. Subsequently on day 7, the cells were then detached using 10 mM EDTA.

### MEFs

MEFs were isolated from E13.5 embryos obtained from pregnant wild-type litter mates, Mavs^-/-^ mice, and STING^gt/gt^ mice following established protocols [90]. Briefly, uterine horns were harvested, and both the outer layer and amniotic sac were carefully removed. Individual embryos were transferred into a 10 cm cell culture dish containing 1× PBS. Heads, tails, and internal organs were excised under sterile conditions, and the remaining embryonic tissue was pooled and finely minced using a No. 22 surgical blade. Tissue fragments were subjected to enzymatic digestion in 3-4 mL of 0.05% trypsin-EDTA and incubated at 37°C for 45 minutes. Following digestion, the tissue was mechanically dissociated by pipetting 20-30 times with a 5 mL serological pipette. The resulting cell suspension was passed through a 100 µm cell strainer to remove undigested tissue. Cells were then pelleted by centrifugation at 1000 × g for 5 minutes, resuspended in MEF isolation medium, and plated into T175 flasks containing 25 mL of medium. Cultures were maintained at 37°C in a humidified 5% CO₂ incubator and allowed to grow to confluency, typically achieved within 3–4 days.

### Mixed glial culture

Mixed glial cell culture was prepared following the method outlined by Guler et al. (2021) [91]. For isolation, P2 pups were decapitated, and their brains were harvested in ice-cold 1X HBSS. The meninges were then carefully removed, and the brain tissue was sliced before being treated with 0.05% DNase I in 1X HBSS (40 units per three pups). The tissue was then gently triturated using a 1 ml and a 200 µl pipette. The resulting mixture was incubated with 0.05% trypsin for 20 minutes at room temperature. After centrifugation, the cell pellet was resuspended in DMEM complete media (2 ml per three pups), gently mixed, and passed through a cell strainer to obtain a single-cell suspension. Following another centrifugation step, the pellet was resuspended in 10 ml of complete media and transferred to a poly-L-lysine-coated T-75 flask. Cells were monitored for growth till 10 days, with media changes occurring on days 1, 2, and 7.

### Cell treatment and virus infection assays

For all virus infection experiments, cells were mock/JEV infected at the indicated MOI for 1 h followed by 1X PBS wash and addition of complete media to the cells. For control experiments, MEFs were transfected with Poly (dA: dT) at 1 mg/ml concentration for 12 h and Poly (I:C) for 6 h using Lipofectamine^TM^ 3000. siRNA treatment was performed using mouse-specific Sting/Tbk1/Caspase-11/Gsdmd/non-targeting (NT) siRNA (30 nM, ON-TARGET plus SMART pool) using the transfection reagent Dharmafect 2 for Neuro2a cells, and Lipofectamine^TM^ RNAimax for MEFs and N9 cells. At 48 h post-transfection, cells were mock/JEV infected at the indicated MOI for 1 h followed by 1X PBS wash and addition of complete media to the cells. For STING inhibitor experiments, H-151 pre-treatment was given by adding DMSO/H-151 (5 µM) for 12 h before virus infection. For STING agonist experiments, virus infection was given at indicated MOI for 1 h, followed by 1XPBS and addition of complete media to the cells. Agonist treatment was given by adding 2 µM diABZI/NFW to mock/JEV infected cells and maintained till the end of experiment. To study TBK1 inhibition, BX-795 treatment was given by adding 250 nM drug/DMSO (control) to mock/JEV-infected cells, which was maintained till the end of the experiment as indicated. For proton channel inhibition experiments, cells were pre-treated with C53 for 2 h, followed by JEV infection or LPS stimulation. At the endpoint of experiment, supernatant was collected to determine virus titer through plaque assay and cytokine levels by ELISA/CBA, cells were then washed and harvested for RNA isolation, western blotting and immunostaining. Every experiment had biological triplicates and was performed two or more times.

### cGAMP production

The DetectX^®^ 2’,3’-cGAMP ELISA Kit was used to quantitatively measure 2’,3’-cGAMP present in lysed cells. Cell lysate collected from mock/UV-JEV/JEV-infected (MOI 2 and 5) and/or untreated/poly (dA:dT) treated MEFs was centrifuged to remove any debris. ELISA was performed according to the manufacturer’s protocol for detecting 2’,3’-cGAMP.

### RNA isolation and real-time quantitative reverse-transcription (qRT) PCR

Total RNA was isolated using Trizol reagent. A total of 1 µg RNA was used to prepare cDNA using ImProm-II™ Reverse Transcription System kit. cDNA was then used to set up qRT-PCR in QuantStudio six-flex RT-PCR machine. Primer sequences were determined from Harvard qPCR primer bank and described in Table S2. JEV RNA level was determined using specific probe and TaqMan reagent. *Gapdh* expression levels were used as an internal housekeeping control. The expression of all other genes was checked using SYBR-green reagent. The fold change in expression of each gene is represented as mean±SD of three or more independent experiments normalized to respective mock/DMSO/NFW/si NT-JEV controls. All experiments had biological triplicates, and qPCR for each sample was performed in technical duplicates.

### Western blotting

Whole cell lysate was using prepared cell lysis buffer (1% Triton X-100 in 50 mM Tris-HCl, 150 mM NaCl, pH 7.5, 1 mM PMSF, and protease inhibitor cocktail; Sigma-Aldrich, Merck) for 1 h at 4°C. Protein concentration was estimated using BCA (bicinchoninic acid) assay kit. Lysates were mixed with 5X Laemmli buffer (40% glycerol, 20% β-mercaptoethanol, 0.04% bromophenol blue, 6% SDS, 0.25 M Tris-HCl pH 6.8) and boiled at 95 °C for 10 min to denature proteins. Equal concentrations of lysates were loaded and separated on SDS-gel followed by transferring on PVDF membrane for immunoblotting. The blots were visualized using Gel Doc XR+ gel documentation system (Bio-Rad) and the intensity of bands was measured using ImageJ (NIH, USA) software. The fold change in protein expression was calculated by normalization to respective loading controls. Data is represented as mean ± SD obtained from three independent experiments.

### Immunostaining, fluorescence microscopy and image processing

For immunofluorescence experiments, cells were seeded on coverslips. JEV infection was given at 5 MOI for 24 h. At the time of harvest, cells were fixed with 2% paraformaldehyde and permeabilized with 0.3% Tween-20 for 30min at RT. This was followed by blocking with 1% Bovine serum albumin in 1X PBS for 1h at RT prior to incubation with primary antibody overnight at 4°C. The cells were washed thrice with 1% BSA for 15 min and then stained using Alexa Fluor labelled specific secondary antibodies for 1h at RT. After washing, mounting of coverslips was done on glass slide ProLong Gold anti-fade reagent with DAPI. Images were acquired on Leica sp8 confocal microscope with 63× (NA 1.4) objective. For co-localization experiments, Z-stacks were acquired at 0.3 µm per slice by sequential scanning with a 63× objective lens. All immunofluorescence experiments were performed in biological duplicates. Images shown are representative of two or more independent experiments

### Subcellular fractionation

Subcellular fractionation was performed following the method outlined by West et al., 2022 [92]. For this, at endpoint of experiment, cells are trypsinised and pelleted down. The pellet is resuspended in 1 ml 1XPBS and then divided into two 1.5 ml tubes labelled as A and B. The tubes are then again centrifuged and the pellet obtained in tube A was used to obtain whole-cell extract (WCE) by resuspending in SDS lysis buffer followed by heating at 95° C for 15 min to lyse the cells. This will serve as a normalization control for the subcellular fractions derived from Tube B. The protocol then employs three detergent-based buffers namely digitonin, NP-40 and SDS lysis buffer and differential centrifugation in a step-wise manner to efficiently and selectively extract cytosolic, mitochondrial, and nuclear fractions respectively. The contents of all the fractions were then denatured and immunoblotted to assess purity.

### ELISA

Culture supernatant collected from mock/JEV-infected and/or untreated/ Poly (da: dt) / Poly (I:C)/LPS treated MEFs or BMDMs was centrifuged to remove any debris. ELISA was performed according to the manufacturer’s protocol for IFNβ, IL6, TNF-α and IL-1β.

### RNA sequencing analysis

RNa sequencing was conducted on biological triplicates of mock-infected control and JEV-infected BMDMs derived from WT and STING^gt/gt^ mice (n = 3). Samples were sequenced on Illumina Platform NovaSeqXplus. An average of ∼100 million 2×150bp read length was sequenced for each sample. Data quality check was performed using FastQC (v0.11.9). The adapter trimming was performed using fastq-mcf program (v1.05) and cutadapt (v4.7). Raw reads were pre-processed to remove adapter and for trimming and quality filtration using Bowtie2 (v2.5.3). The paired-end reads were aligned to the reference Mouse genome downloaded from the Ensembl database (genome-build GRCm38.p6; genome-version GRCm38; genome-build-accession NCBI: GCA_000001635.8). Alignment was performed using HISAT2 (v2.2.1) aligner. Reads mapping to ribosomal and mitochondrial genome were removed before performing alignment. Also, RNA-Seq detailed quality control was performed using RNA-SeQC (v2.4.2), RSeQC (v3.0.1) and MultiQC (v1.21).The raw read counts were estimated using featureCounts (v2.0.6). Read counts were normalized using DESeq2 to get the normalized counts. Additionally, the aligned reads were used for estimating expression of the genes using cuflinks (v2.2.1). The expression values were reported in FPKM (Fragments per kilobase per million) units for each gene. To gain insight into the nature of DEGs uniquely expressed in WT and STING^gt/gt^ during JEV infection multiple comparisons were considered with significant differential expression cutoff: P value < 0.05 & Fold change >=1.5 or <=-1.5. Over Representation Analysis (ORA), Gene Set Enrichment Analysis (GSEA) and Pathway analysis was performed for significantly differentially expressed up and down regulated protein coding genes. ORA and GSEA was performed for gene ontology, disease and pathway analysis. DAVID (DAVID v6.8) (The Database for Annotation, Visualization and Integrated Discovery) and KEGG were used for pathway analysis. ORA and GSEA analysis were performed using an R Bioconductor package clusterProfiler (v3.14.3).

### Cytokines bead array (CBA)

WT and STING^gt/gt^ BMDMs (biological triplicates) were mock/JEV infected at 5 MOI for 1 h. At 24 hpi, complete media was added to the cells for 24h. Supernatants were collected to analyse cytokine levels of IL-6, TNF-α, MCP-1, RANTES, and IL-1β. The cytokines were quantified using LEGENDPLEX MU anti-virus response panel 6 plex assay kit following the manufacturer’s instructions. Data analysis was performed using LEGENDplex™ Multiplex Assay Software, and cytokine concentrations were determined based on their respective standard curves. All CBA assays were performed two or more times, and representative data from one experiment is shown.

### Animal experiments

The mouse-adapted strain JEV-S3 was generated in 3–4-day-old C57BL/6 mouse pups, following the protocol described by Tripathi et al. (2021) [54]. In brief, the pups were intracranially infected with JEV (10⁵ PFU). By 3–4 days post-infection, symptoms of JEV infection, including movement impairment and persistent shivering or body tremors were observed. The pups were then euthanized, and their brain tissues were harvested, homogenized in incomplete MEM medium, and the supernatant containing the infectious virus was quantified through plaque assay. For survival, body weight and other experiments, 3-to 4-week-old C57BL/6 and STING^gt/gt^ mice were weighed followed by their grouping into mock/JEV by matching weight. Mice were intra-peritoneally infected with 10^7^ PFU JEV, and equal volume of incomplete MEM was injected intra-peritoneally into mock-infected group. Mice were then monitored for visual clinical symptoms of viral encephalitis like reduction in body weight, piloerection, tremor, loss in movement, paralysis and mortality. For survival and body weight study, mice were followed up till 15 days post infection. For other assays, mice were sacrificed 24 h post onset of symptoms and brain of the mice was harvested. The brain tissue was then homogenized to determine JEV RNA levels and titer. To determine cytokine and protein levels of inflammatory markers, brain tissue was homogenized in lysis buffer and 30 µg of protein was used for each assay. The cytokines were quantified using LEGENDPLEX MU anti-virus response panel 6 plex assay kit following the manufacturer’s instructions. Data analysis was performed using LEGENDplex™ Multiplex Assay Software, and cytokine concentrations were determined based on their respective standard curves.

### Cytotoxicity assay

Cells were treated with DMSO/NFW/H-151/BX-795/diABZI/C53 at indicated concentrations for 24 h. The MTT assay was performed as per the product manual to study the effect of H-151/BX-795/diABZI/C53 on cell viability normalized to respective control.

## Statistical analysis

Statistical analysis was performed using paired/unpaired Student’s t-test and Dunnett test/two-way ANNOVA followed by Sidak’s multiple comparisons test and Log-rank (Mantel-Cox) test. Differences were considered significant at P values of *p < 0.05, **p < 0.01, ***p < 0.001, and ****p < 0.0001, as indicated in the figure legends. Error bar indicates means ± SD/SEM. All graphs were plotted and analysed using GraphPad Prism 8.

## Data availability

The datasets generated in this study are available in public repositories: Indian Biological Data Centre (IBDC) Study accession INRP000339, INSDC Project Accession: PRJEB90219 / ERP173232.

## Funding Information

This work was supported by ANRF (SERB) grant CRG/2021/006523 to MK. SC, DP, and V.S were supported by DBT fellowships.

## Acknowledgements

All Virology lab members are acknowledged for useful discussions and constant support. We acknowledge Dr. Shailendra Chauhan for technical help with RNA-sequencing data analysis. We thank the facilities and staff of Advanced Technology Platform Centre, Small Animal Facility and Indian Biological Data Centre. We are also thankful to Mr. Gaurav Lukhar for his help with animal experiments. The schematic model was created using bioRender.

## Disclosure Statement

The authors have no conflict of interest to declare.

## Notes

### Competing Interest Statement

The authors have declared no competing interest.

