## Supplementary Data for "STING is a key driver of Japanese encephalitis virus induced inflammatory response"

**Supplementary Data File**

Supplementary Figures S1-S10

Supplementary Tables S1-S2

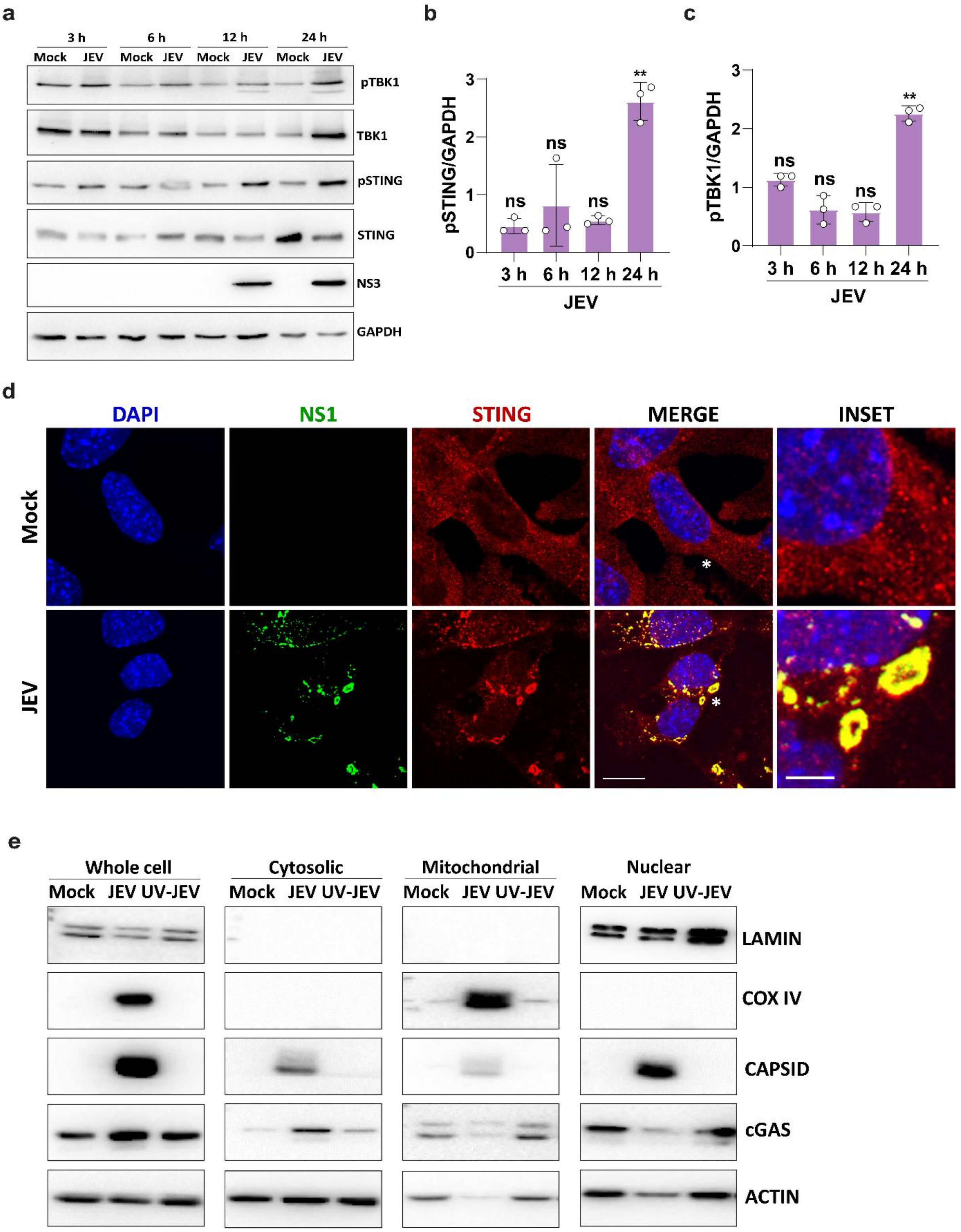


**Figure S1: JEV infection activates STING and cGAS**

MEFs were mock/JEV infected at 2 MOI for 3, 6, 12 and 24 h. (a) Protein lysates were immunoblotted with pSTING, STING, pTBK1, TBK1, JEV-NS3 and GAPDH (loading control) antibodies. (b,c) BAR graph shows the normalized pSTING and pTBK1 protein expression

(JEV/mock) at indicated timepoints for three independent experiments. (d) MEFs were mock/JEV infected with JEV (5 MOI, 24h). Cells were stained with JEV-NS1 (green) and STING (red) antibodies. Nucleus was stained with DAPI. Images were acquired on a confocal microscope using a 63x objective. Scale bar 10µm. Images are representative of two independent experiments. (e) MEFs were mock/JEV/UV-JEV infected at 2 MOI for 24 h followed by subcellular fractionation to isolate whole cell, cytosolic, nuclear and mitochondrial fractions and analysed by western blotting with LAMIN A/C, COX IV, CAPSID, cGAS and ACTIN antibodies.

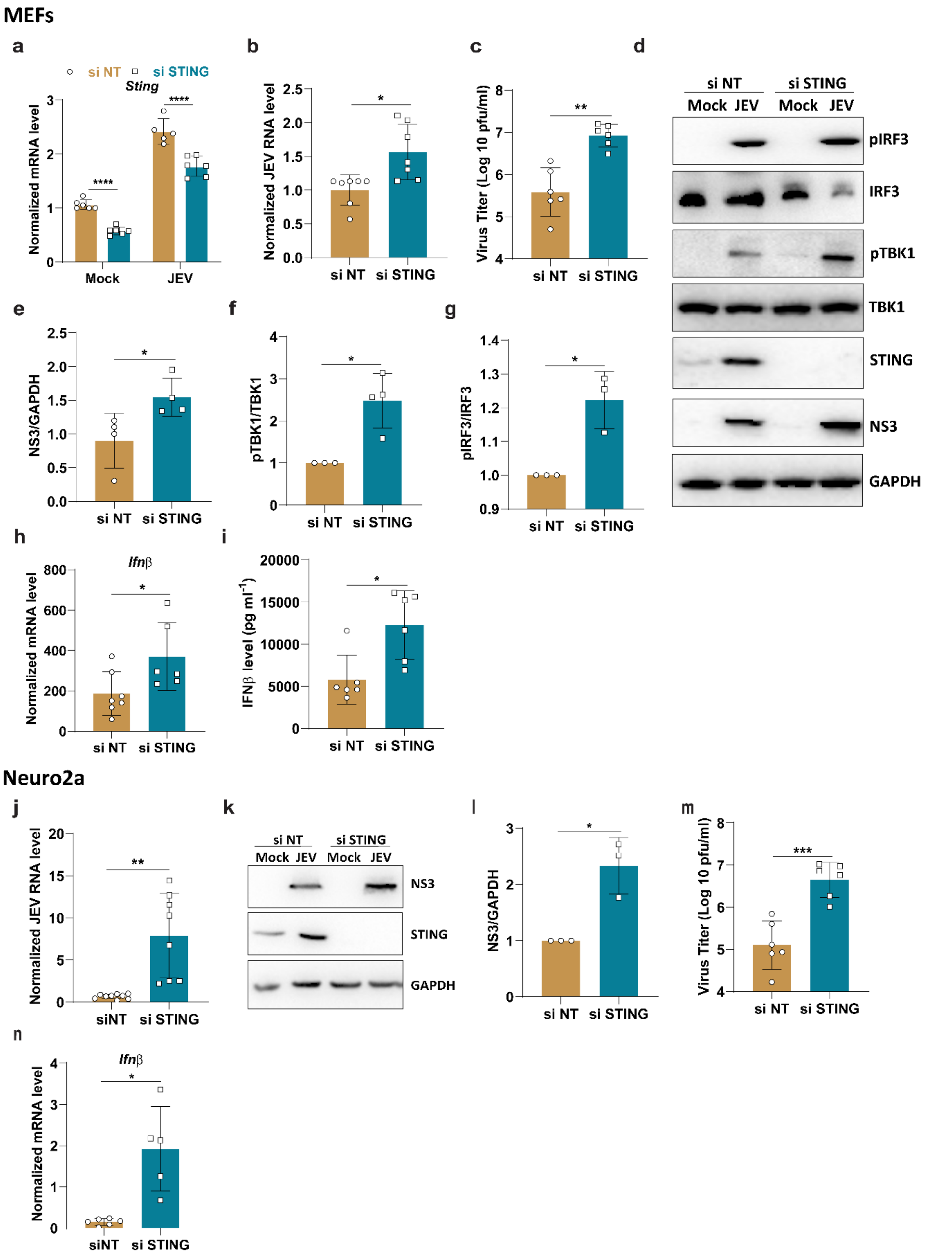


**Figure S2: STING depletion enhances virus replication and IFN production**

NT/STING siRNA (30 nM, 48 h) treated MEFs were mock/JEV infected at 2 MOI for 24 h. Total RNA was isolated to quantitate *Sting* mRNA expression (a) and JEV-RNA levels (b) by qRT-PCR.

(c) Culture supernatant was collected to determine extracellular virus titer by plaque assay. (d) Protein lysates were immunoblotted with pIRF3, IRF3, pTBK1, TBK1, STING, JEV-NS3 and GAPDH (loading control) antibodies. Bar graph shows the normalized JEV-NS3 (e), pTBK1/TBK1 (f) and pIRF3/IRF3 (g) protein expression (JEV/mock). (h) *Ifnβ* transcript levels were quantitated by qRT-PCR and normalized to si NT-Mock. (i) Culture supernatant collected from mock/JEV-infected MEFs was used to perform ELISA for IFNβ. NT/STING siRNA (30 nM, 48 h) treated N2a cells were mock/JEV infected at 5 MOI for 24 h. Total RNA was isolated to quantitate JEV-RNA levels (j) by qRT-PCR. (k) Protein lysates were immunoblotted with JEV-NS3, STING and GAPDH (loading control) antibodies. (l) Bar graph shows the normalized JEV-NS3 protein expression ratio (JEV/mock). (m) Culture supernatant was collected to determine extracellular virus titer by plaque assay. (n) *Ifnβ* transcript levels were quantitated by qRT-PCR and normalized to si NT-Mock.

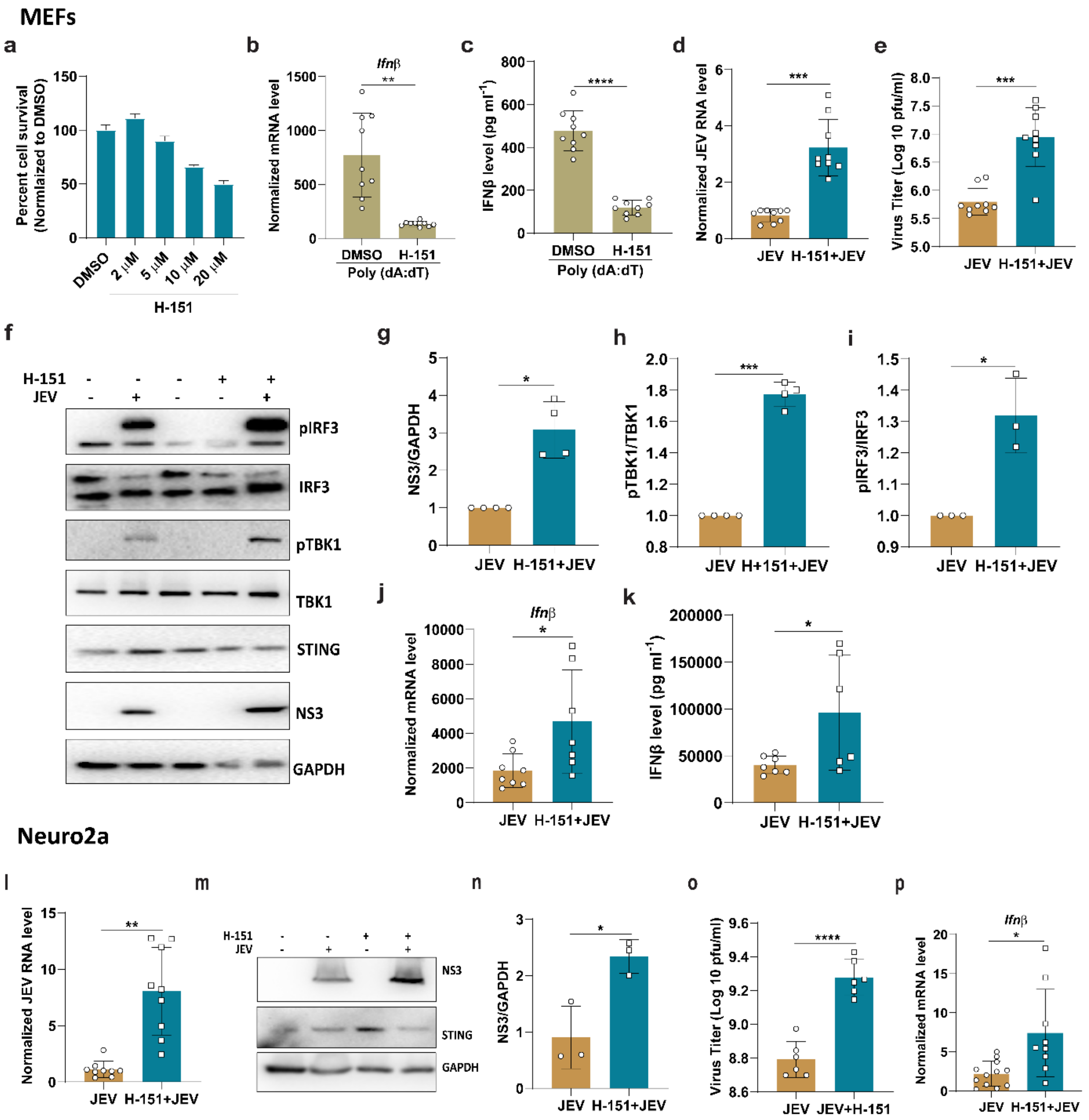


**Figure S3: Pharmacological inhibition of STING enhances virus replication**

MEFs were treated with DMSO or 2/5/10/20 µM H-151 for24 h. (a) Cell viability was studied using the MTT assay. The bar graph shows % cell viability measured relative to DMSO-treated cells. MEFs were pretreated with DMSO/ H-151 (5 µM) for 12 h followed by transfection with Poly (dA: dT) (1 mg/ml) for 12 h. Total RNA was isolated to quantitate transcript levels of *Ifnβ* (b). (c) Culture supernatant was collected to quantify IFNβ cytokine levels. MEFs were pretreated with DMSO or 5 µM of H-151 for 12h followed by mock/JEV infection at 2 MOI for 24 h. (d) Total RNA was isolated to quantitate JEV RNA levels by qRT-PCR. (e) Culture supernatant was collected to quantify extracellular virus titre by plaque assay. (f) Protein lysates were prepared and analyzed through western blotting with pTBK1, TBK1, pIRF3, IRF3, JEV-NS3, STING and GAPDH (loading control) antibodies. Bar graph shows the normalized NS3/GAPDH (g) , pTBK1/TBK1 (h) and pIRF3/IRF3 (i) protein expression ratio (JEV/mock) for three independent experiments. (j) *Ifnβ* transcript levels were quantitated by qRT-PCR (k) Culture supernatant collected from mock/JEV-infected MEFs was used to perform ELISA for IFNβ. N2a cells were pretreated with DMSO or 5 μM of H-151 for 12h followed by mock/JEV infection at 5 MOI for 24 h. (l) Total RNA was isolated to quantitate JEV RNA levels by qRT-PCR. (m) Protein lysates were prepared and analyzed through western blotting with JEV-NS3, STING and GAPDH (loading control) antibodies. (n) Bar graph shows the normalized NS3/GAPDH protein expression ratio (JEV/mock) for three independent experiments. (o) Culture supernatant was collected to quantify extracellular virus titre by plaque assay. (p) *Ifnβ* transcript levels were quantitated by qRT-PCR. Data presented is mean ± SD of values obtained from 3 independent experiments. Student t-test was used to calculate P values. *P < 0.05, **P < 0.01, ****P < 0.0001. ns, non-significant.

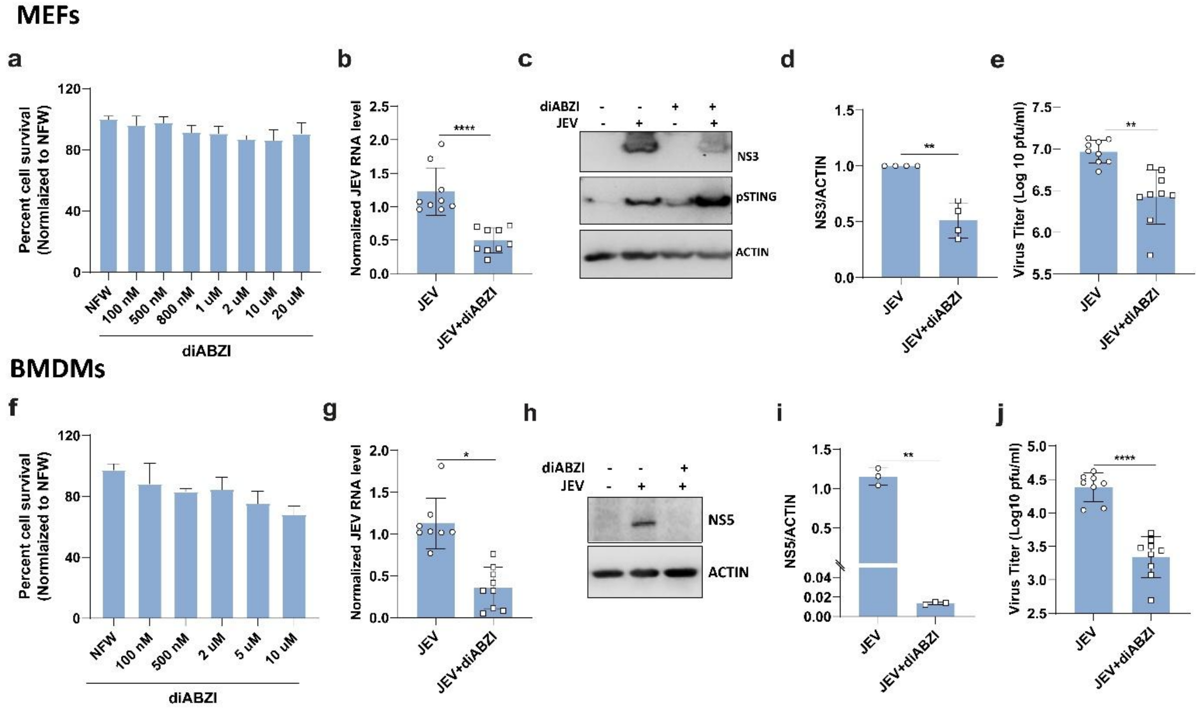


**Figure S4: Pharmacological activation of STING restricts virus replication**

MEFs were treated with DMSO or 100/500/800 nM and 1/2/10/20 µM diABZI for24 h. (a) Cell viability was studied using the MTT assay. The bar graph shows % cell viability measured relative to NFW-treated cells. MEFs were mock/JEV infected followed by treatment with 2 µM diABZI and at 24hpi, cells were harvested. Total RNA was isolated and JEV-RNA levels were quantitated by qRT-PCR (b). (c) Protein lysates were immunoblotted with JEV-NS3, pSTING and ACTIN (loading control) antibodies. (d) Bar graph shows the normalized JEV-NS3 protein expression (JEV/mock). (e) Culture supernatant was collected to determine extracellular virus titer by plaque assay. BMDMs were treated with NFW or 100/500 nM and 2/5/10 µM diABZI for24 h. (f) Cell viability was studied using the MTT assay. The bar graph shows % cell viability measured relative to NFW-treated cells. BMDMs were mock/JEV infected at 5 MOI followed by treatment with 2 µM diABZI and at 24hpi cells were harvested. Total RNA was isolated to determine JEV-RNA levels (g) by qRT-PCR. (h) Protein lysates were immunoblotted with JEV-NS5 and ACTIN (loading control) antibodies. (i) Bar graph shows the normalized JEV-NS5 protein expression (JEV/mock). (j) Culture supernatant was collected to determine extracellular virus titer by plaque assay.

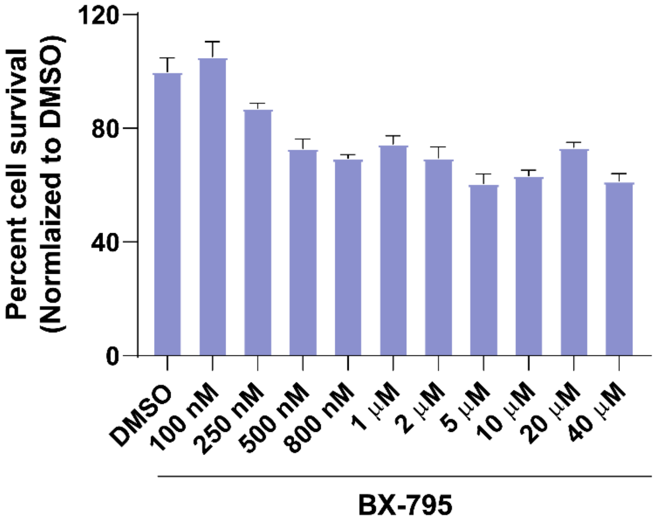


**Figure S5: Cell viability assay for BX-795**

MEFs were treated with DMSO or 100/250/500/800 nM and 1/2/5/10/20/40 µM BX-795 for 24 h. (a) Cell viability was studied using the MTT assay. The bar graph shows % cell viability measured relative to DMSO-treated cells.

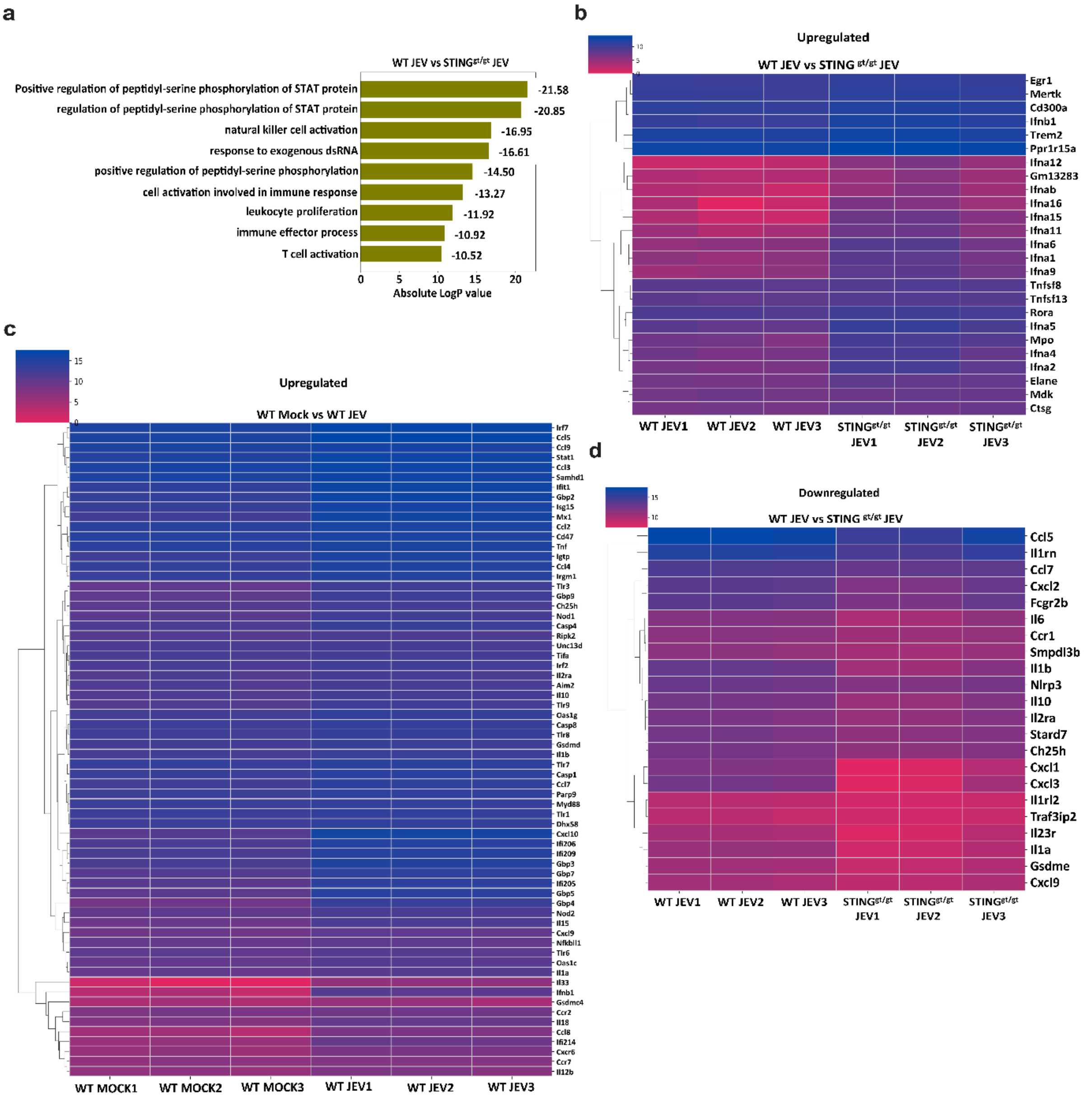


**Figure S6: Transcriptome analysis of JEV infected WT and STING$^{gt/gt}$ BMDMs**

(a) Pathway enrichment analysis of the upregulated DEGs in JEV-infected STING$^{gt/gt}$ BMDMs compared to WT BMDMs. (b) Heatmap showing log2-transformed gene expression levels for selected upregulated genes across different samples. The color scale uses a cerise (bright pinkish-red) to studio (blue) palette. Genes are clustered hierarchically based on expression similarity, while samples are arranged with WT JEV samples first (columns 1-3) followed by STING$^{gt/gt}$ JEV samples (columns 4-6). The dendrogram on the left shows gene clustering patterns. (c) Heatmap showing log2-transformed gene expression levels for selected upregulated genes across different samples. The color scale uses a cerise (bright pinkish-red) to studio blue palette. Genes are clustered hierarchically based on expression similarity, while samples are arranged with WT Mock samples first (columns 1-3) followed by WT JEV samples (columns 4-6). The dendrogram on the left shows gene clustering patterns. (d) Heatmap showing log2-transformed gene expression levels for selected downregulated genes across different samples. The color scale uses a cerise (bright pinkish-red) to studio (blue) palette. Genes are clustered hierarchically based on expression similarity, while samples are arranged with WT JEV samples first (columns 1-3) followed by STING$^{gt/gt}$ JEV samples (columns 4-6). The dendrogram on the left shows gene clustering patterns.

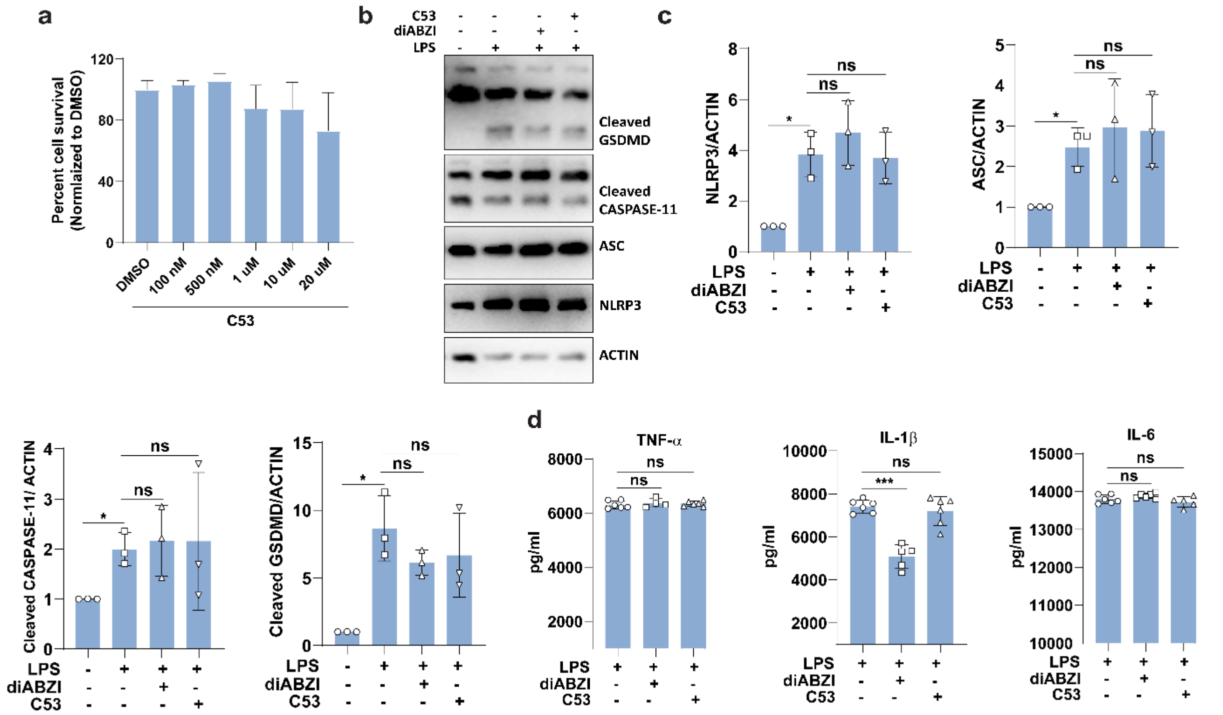


**Figure S7: STING proton channel inhibition does not impact LPS induced inflammatory response**

BMDMs were treated with DMSO or 100/500 nM and 1/10/20 µM C53 for24 h. (a) Cell viability was studied using the MTT assay. The bar graph shows % viability measured relative to DMSO-treated cells. (b) BMDMs were either pre-treated with C53 (10 µM) for 2 h, then stimulated with LPS (1 µg/ml) for 4 h and nigericin (1 µM) for 1 h, or co-treated with diABZI (2 µM) and LPS (1 µg/ml) for 4 h, followed by nigericin (1 µM) for 1 h. Protein lysates were prepared and immunoblotted with GSDMD, CASPASE-11, ASC, NLRP3, and ACTIN (loading control) antibodies. (c) Bar graph shows the normalized NLRP3, ASC, cleaved CASPASE-11 and cleaved GSDMD protein expression (LPS/mock). (d) Culture supernatant was collected to determine the extracellular release of TNF-α, IL-1β and IL-6 and from mock/LPS treated BMDMs through ELISA.

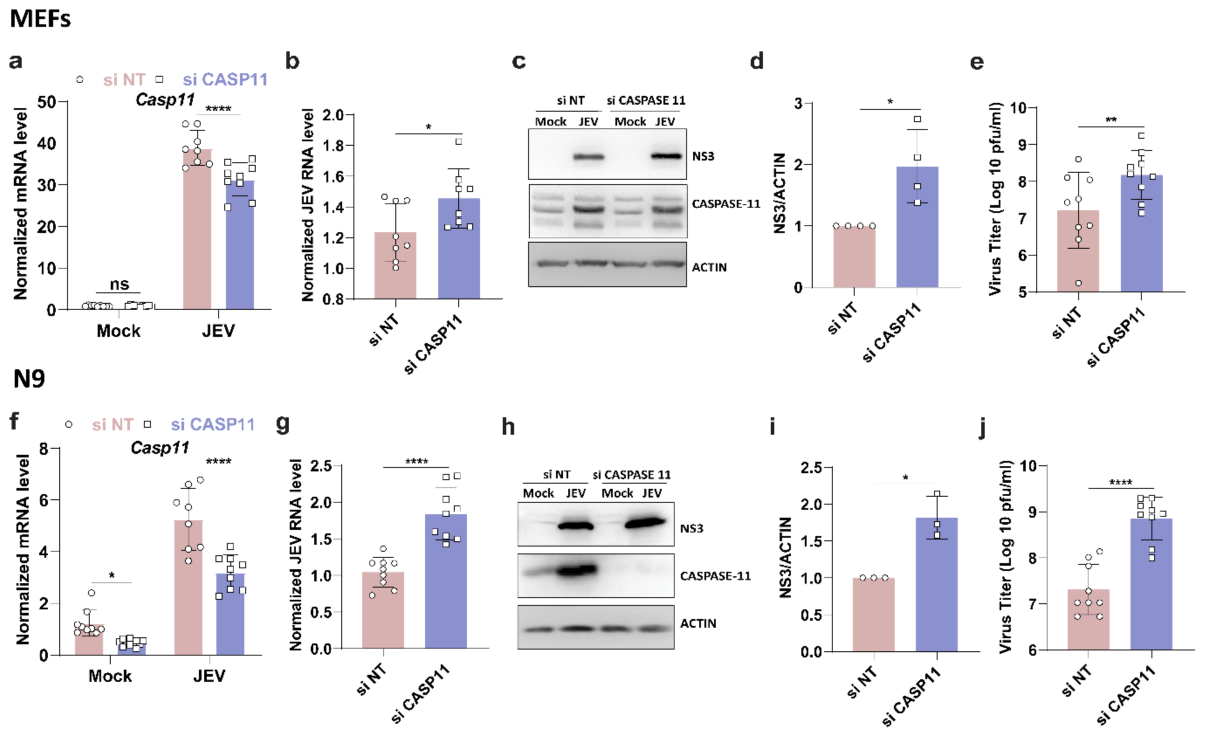


**Figure S8: CASPASE-11 restricts virus replication in fibroblast and microglial cells**

NT/CASPASE11 siRNA (30 nM, 48 h) treated MEFs were mock/JEV infected at 2 MOI for 24 h. Total RNA was isolated to quantitate *Caspase11* mRNA expression (a) and JEV-RNA levels (b) by qRT-PCR. (c) Protein lysates were immunoblotted with CASPASE 11, JEV-NS3 and ACTIN (loading control) antibodies. (d) Bar graph shows the normalized JEV-NS3 protein expression (JEV/mock). (e) Culture supernatant was collected to determine extracellular virus titer by plaque assay. NT/CASPASE11 siRNA (30 nM, 48 h) treated N9 cells were mock/JEV infected at 2 MOI for 24 h. Total RNA was isolated to quantitate *Caspase-11* mRNA expression (f) and JEV-RNA levels (g) by qRT-PCR. (h) Protein lysates were immunoblotted with JEV-NS3, CASPASE-11and ACTIN (loading control) antibodies. (i) Bar graph shows the normalized JEV-NS3 protein expression (JEV/mock). (j) Culture supernatant was collected to determine extracellular virus titer by plaque assay.

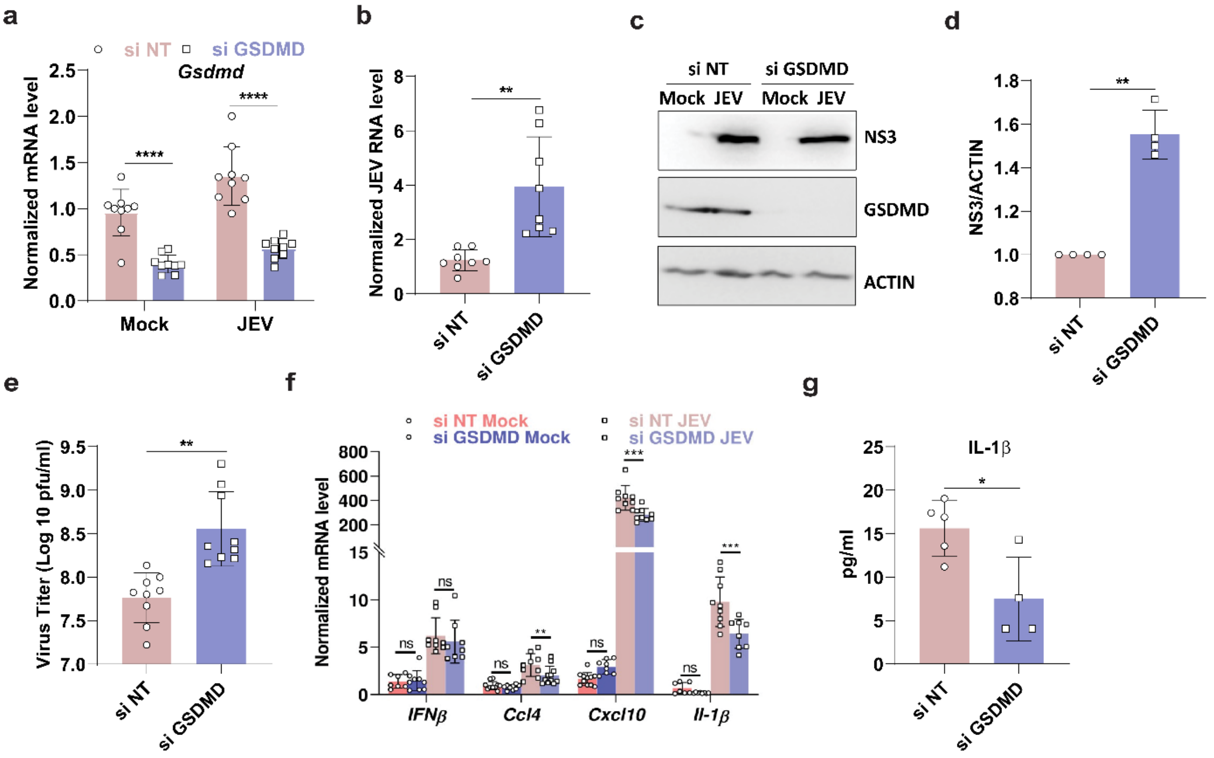


**Figure S9: GSDMD plays an antiviral role in microglial cells**

NT/GSDMD siRNA (30 nM, 48 h) treated N9 cells were mock/JEV infected at 2 MOI for 24 h. Total RNA was isolated to quantitate *Gsdmd* mRNA expression (f) and JEV-RNA levels (g) by qRT-PCR. (h) Protein lysates were immunoblotted with JEV-NS3, GSDMD and ACTIN (loading control) antibodies. (i) Bar graph shows the normalized JEV-NS3 protein expression (JEV/mock). (j) Culture supernatant was collected to determine extracellular virus titer by plaque assay. (k) Transcript levels of *Ifnβ, Ccl4, Cxcl10* and *Il-1β* were determined by qRT-PCR. (l) Culture supernatant collected from mock/JEV-infected N9 cells was used to perform ELISA for IL-1β.

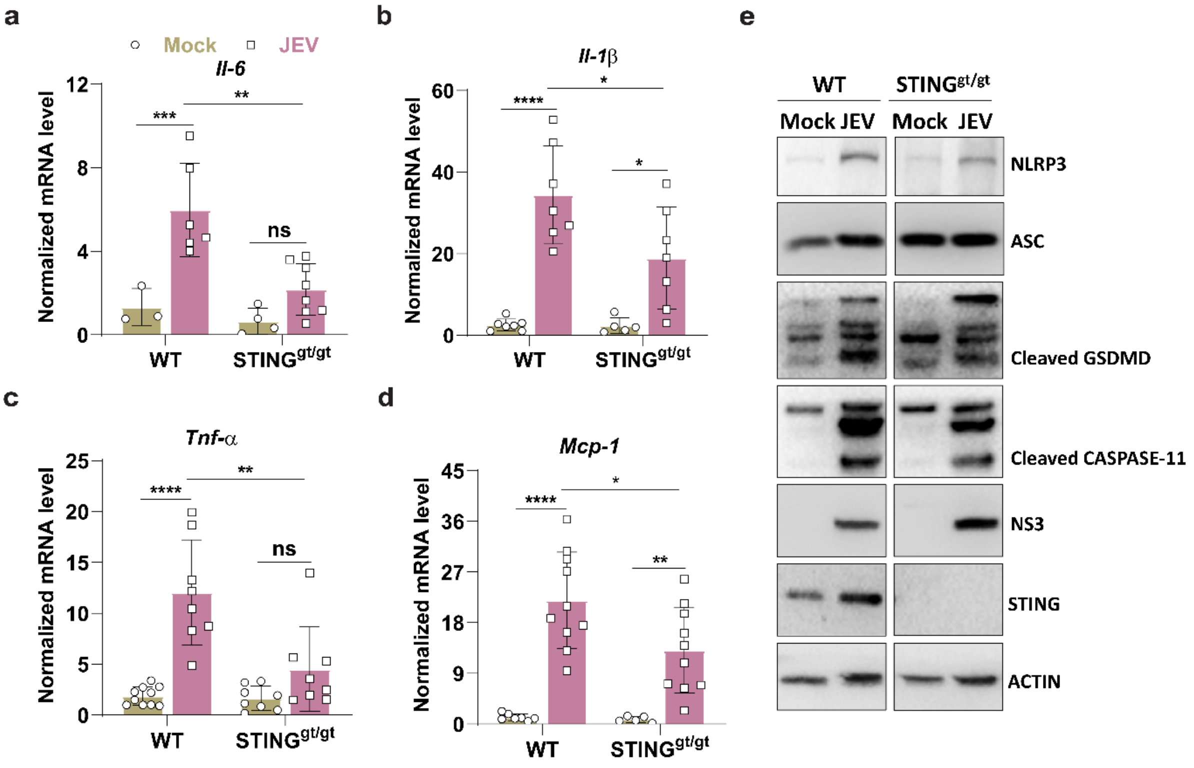


**Figure S10: STING deficiency suppressed JEV-induced non-canonical inflammasome activation and pyroptotic cell death.**

Mixed glial cells isolated form brain of 2 days old WT and STING$^{gt/gt}$ C57BL/6 pups were either mock/JEV infected at 5 MOI for 24 h (a,b,c,d) Total RNA was isolated to transcript levels of *Il-6, Il-1β, Tnf-α* and *Mcp-1* were determined using qRT-PCR. (e) Protein lysates were immunoblotted with NLRP3, ASC, GSDMD, CASPASE-11, JEV-NS3, STING and ACTIN (loading control) antibodies. Data presented is mean ± SD of values obtained from 3 independent experiments. Student t-test was used to calculate P values. *P < 0.05, **P < 0.01, ****P < 0.0001. ns, non-significant.

### Table S1: Chemical reagents used in this study

| S. No. | Name of Reagent | Catalogue No. |
|---|---|---|
| 1. | Agarose Type VII | Sigma-Aldrich (A4018) |
| 2. | BCA assay kit | G-Biosciences (786-570) |
| 3. | Bromophenol blue | Sigma-Aldrich (B0126-25G) |
| 4. | BX795 | Medchem Express (HY-10514) |
| 5. | Compound 53 (STING agonist-12) | Medchem Express (HY-147010) |
| 6. | Crystal violet | SRL (28376) |
| 7. | Deoxyribonuclease I (DNase I) | SRL (61824) |
| 8. | DetectX® 2′,3′-Cyclic GAMP ELISA Kit. | Arbor assays (K067-H1/H5) |
| 9. | diABZI (compound 3) | InvivoGen (tlrl-diabzi-2) |
| 10. | Dimethyl sulfoxide (DMSO) | Sigma-Aldrich (276855-250ML) |
| 11. | DL-Dithiothreitol (DTT) | SRL (3483-12-3) |
| 12. | Dimethyl sulfoxide (DMSO) | Sigma-Aldrich (322415-100ML) |
| 13. | EDTA | Sigma-Aldrich (E9884-500G) |
| 14. | Formaldehyde | Merck (D12F721465) |
| 15. | H-151 | Merck (SML2437-5MG) |
| 16. | HBSS | Gibco (14175095) |
| 17. | HEPES | Sigma-Aldrich (H3375-25G) |
| 18. | ImProm-II™ Reverse Transcription System | Promega (A3800) |
| 19. | LPS | Sigma-Aldrich (L2630) |
| 20. | MTT 3-(4,5-Dimethylthiazol-2-yl)-2,5-Diphenyltetrazolium Bromide | VWR life science, (0793-1G) |
| 21. | Nigericin | InvivoGen (tlrl-nig) |

| | | |
|---|---|---|
| 22. | Phenylmethylsulfonyl fluoride (PMSF) | Sigma-Aldrich (329-98-6) |
| 23. | Premix Ex Taq™ (Probe qPCR) | Takara (RR390A) |
| 24. | ProLong™ Gold Antifade Mountant with DAPI | Invitrogen (P36935) |
| 25. | Protease inhibitor cocktail (PI) | Sigma-Aldrich (P8340) |
| 26. | Puromycin | InvivoGen (ant-pr-1) |
| 27. | PVDF membrane | Merck Millipore (IPVH00010) |
| 28. | Random hexamer | Sigma-Aldrich (H0268) |
| 29. | RBC lysis buffer | GCC Biotech (19114B1076) |
| 30. | Recombinant RNasin® Ribonuclease inhibitor | Promega (Ref N2515) |
| 31. | SDS | Sigma-Aldrich (L3771-500G) |
| 32. | SYBR® Premix Ex Taq™ | Takara (RR420A) |
| 33. | Triton™ X-100 | Sigma-Aldrich (T9284-500ML) |
| 34. | Trizol reagent (RNAiso Plus) | Takara (9109) |
| 35. | Tween 20 | G-Biosciences (RC1227) |
| | **Media & other additives** | **Catalogue No.** |
| 36. | 2XMEM | Himedia (AL178A-500ML) |
| 37. | DMEM, powder, high glucose | Gibco (12100-046) |
| 38. | FBS- QUALIFIED, 500ML | Gibco (10270106) |
| 39. | LEIBOVITZ L 15 MED, 1X10L | Gibco (41300070) |
| 40. | L929 conditioned media | Culture supernatant of L929 fibroblast cells (After 6-8 days starvation of L929 cells) |
| 41. | L-Glutamine | Himedia (TCL012) |
| 42. | MEM EARLES, 10X1L | Gibco (61100061) |
| 43. | MEM Amino Acids Solution (50X) | Gibco (11130051) |
| 44. | Penicillin-Streptomycin | Himedia (A007-100ML) |

| | | |
|---|---|---|
| 45. | RPMI 1640, 1X50L | Gibco (31800105) |
| 46. | Sodium pyruvate | Himedia (TCL015-100ML) |
| 47. | Trypsin - EDTA Solution | Himedia (TCL007) |
| | **Antibodies** | **Catalogue No.** |
| 48. | Alexa fluor$^{TM}$ 488 goat anti-rat IgG (H+L) | Invitrogen (940882) |
| 49. | Alexa fluor$^{TM}$ 568 goat anti-rabbit IgG (H+L) | Invitrogen (A-11011) |
| 50. | Alexa fluor$^{TM}$ 647 donkey anti-mouse IgG (H+L) | Invitrogen (A-21463) |
| 51. | ACTIN | CST (4970S) |
| 52. | ASC | CST (67824S) |
| 53. | ATG5 | CST (12994S) |
| 54. | BETA-TUBULIN | Abcam (ab21058) |
| 55. | CASPASE-11 | Abcam (ab180673) |
| 56. | cGAS | CST (31659S) |
| 57. | COX-IV | CST (11967S) |
| 58. | GAPDH | GeneTex (GTX100118) |
| 59. | GSDMD | Abcam (ab209845) |
| 60. | IKKε | CST (3416S) |
| 61. | IRF3 | CST (4302S) |
| 62. | JEV-CORE | GeneTex (GTX131368) |
| 63. | JEV-NS1 | Abcam (ab41651) |
| 64. | JEV-NS1 | GeneTex (GTX633820) |
| 65. | JEV-NS3 | GeneTex (GTX125868) |
| 66. | JEV-NS5 | GeneTex (GTX131359) |
| 67. | NF-κβ | CST (8242S) |
| 68. | NLRP3 | R&D (MAB7578) |

| | | |
|---|---|---|
| 69. | NLRP3 | CST (15101S) |
| 70. | pIKKε | CST (8766S) |
| 71. | pIRF3 | CST (4947S) |
| 72. | pNF-κβ | CST (3033S) |
| 73. | pSTING | PA5-105674 |
| 74. | pSTING | CST (72971S) |
| 75. | pTBK1 | CST (5483S) |
| 76. | Peroxidase AffiniPure Donkey Anti-Mouse IgG (H+L) | Jackson ImmunoResearch (715-035-150) |
| 77. | Peroxidase AffiniPure Donkey Anti-Rabbit IgG (H+L) | Jackson ImmunoResearch (711-035-152) |
| 78. | STING | CST (13647S) |
| 79. | TBK1 | CST (38066S) |
| | **siRNA/Transfection reagents** | **Catalogue No.** |
| 80. | DharmaFECT 2 | Dharmacon (T-2002-02) |
| 81. | Lipofectamine™ RNAimax | Invitrogen (13778030) |
| 82. | Lipofectamine™ 3000 | Invitrogen (L3000015) |
| 83. | ON-TARGETplus Non-targeting (NT) | Dharmacon (D-001810-10-20) |
| 84. | ON-TARGETplus Mouse si cgas | Sigma-Aldrich (SASI_MM01_00129826) |
| 85. | ON-TARGETplus Mouse si Sting | Sigma-Aldrich (SASI_MM02_00424289) |
| 86. | ON-TARGETplus Mouse si TBK1 | Dharmacon (L-063162-00-0005) |
| 87. | ON-TARGETplus Mouse si CASPASE-11 | Sigma-Aldrich (SASI_Mm01_00176950) |
| 88. | ON-TARGETplus Mouse si GSDMD | Dharmacon (M-046650-01-0005) |
| | **Cytokines** | **Catalogue No.** |
| 89. | LEGENDPLEX MU Anti-Virus Response Panel (13-plex) | Biolegend (740622) |
| 90. | Mouse IFN-β Flex Set | BD Bioscience (740145) |

| | | |
|---|---|---|
| 91. | Mouse IL-1β Flex Set | BD Bioscience (740634) |
| 92. | Mouse IL-6 Flex Set | BD Bioscience (740629) |
| 93. | Mouse MCP-1 Flex Set | BD Bioscience (740627) |
| 94. | Mouse RANTES Flex Set | BD Bioscience (740628) |
| 95. | Mouse TNF-α Flex Set | BD Bioscience (741260) |
| 96. | Mouse/Rat Soluble Protein Master Buffer Kit | BD Bioscience (558266) |
| | **ELISA kits** | **Catalogue No.** |
| 97. | IFN-β MOUSE IFN-BETA DUOSET ELISA, 5 PLATE 1 KIT | R&D Systems (DY410-05) |
| 98. | MOUSE IL-1 BETA/IL-1F2 DUOSET ELISA, 5 PLATE 1 KIT | R&D Systems (DY406-05) |
| 99. | MOUSE IL-6 DUOSET ELISA, 5 PLATE 1 KIT | R&D Systems (DY401-05) |
| 100. | MOUSE TNF-ALPHA DUOSET ELISA, 5 PLATE 1 KIT | R&D Systems (DY8234- 05) |
| 101. | DUOSET ELISA ANCILLARY REAGENT KIT 2 1 KIT | R&D Systems (DY008B) |
| 102. | SUBSTRATE REAGENT PACK 1 PACK | R&D Systems (DY999B) |
| | **Animal Models** | |
| 104. | C57BL/6J | The Jackson Laboratory (JAX stock # 000664) |
| 105. | C57BL/6J-*Sting1*$^{gt}$/J | The Jackson Laboratory (JAX stock # 017537) |
| 106. | B6;129-*Mavs*$^{tm1Zjc}$/J | The Jackson Laboratory (JAX stock # 008634) |
| 107. | AG129 | Marshall BioResources |

**Table S2: Primers used in this study**

| S.No. | Genes | Forward (5'-3') sequence | Reverse (5'-3') sequence |
|---|---|---|---|
| 1. | Caspase-11 | ACAAACACCCTGACAAACCAC | CACTGCGTTCAGCATTGTTAAA |
| 2. | Ccl4 | TTCCTGCTGTTTCTCTTACACCT | CTGTCTGCCTCTTTTGGTCAG |
| 3. | cgas | GAGGCGCGGAAAGTCGTAA | TTGTCCGGTTCCTTCCTGGA |
| 4. | Cxcl10 | CCAAGTGCTGCCGTCATTTTC | CCCTATGGCCCTCATTCTCA |
| 5. | Gapdh | CGTCCCGTAGACAAAATGGT | TTGATGGCAACAATCTCCAC |
| 6. | Gsdmd | CCATCGGCCTTTGAGAAAGTG | ACACATGAATAACGGGGTTTCC |
| 7. | Ifn-β | AAGAGTTACACTGCCTTTGCCATC | CACTGTCTGCTGGTGGAGTTCATC |
| 8. | Il-1β | TCCAAGAAAGGACGAACATTCG | TGAGGACATCTCCCACGTCAA |
| 9. | Il-6 | CTGCAAGAGACTTCCATCCAG | AGTGGTATAGACAGGTCTGTTGG |
| 10. | Irf7 | GAGACTGGCTATTGGGGGAG | GACCGAAATGCTTCCAGGG |
| 11. | JEV | AGAGCACCAAGGGAATGAAATAGT<br>Taqman probe: CCACGCCACTCGACCCATAGACTG (5′ end FAM, 3′end TAMRA). | AATAAGTTGTAGTTGGGCACTCTG |
| 12. | Mavs | CTGCCTCACAGCTAGTGACC | CCGGCGCTGGAGATTATTG |
| 13. | Mcp-1 | CAAGAAGGAATGGGTCCAGA | GCTGAAGACCTTAGGGCAGA |
| 14. | Sting | CTACATTGGGTACTTGCGGTT | GCACCACTGAGCATGTTGTTATG |
| 15. | Tbk1 | ACTGGTGATCTCTATGCTGTCA | TTCTGGAAGTCCATACGCATTG |
| 16. | Tnf-α | CCCTCACACTCAGATCATCTTCT | GCTACGACGTGGGCTACAG |
